# Rewiring an olfactory circuit by altering the combinatorial code of cell-surface proteins

**DOI:** 10.1101/2025.03.01.640986

**Authors:** Cheng Lyu, Zhuoran Li, Chuanyun Xu, Jordan Kalai, Liqun Luo

**Affiliations:** Department of Biology and Howard Hughes Medical Institute, Stanford University, Stanford, CA 94305, USA; Biology Graduate Program, Stanford University, Stanford, CA 94305, USA

## Abstract

Proper brain function requires the precise assembly of neural circuits during development. Despite the identification of many cell-surface proteins (CSPs) that help guide axons to their targets^1,2^, it remains largely unknown how multiple CSPs work together to assemble a functional circuit. Here, we used synaptic partner matching in the *Drosophila* olfactory circuit^3,4^ to address this question. By systematically altering the combination of differentially expressed CSPs in a single olfactory receptor neuron (ORN) type, which senses a male pheromone that inhibits male-male courtship, we switched its connection from its endogenous postsynaptic projection neuron (PN) type nearly completely to a new PN type that promotes courtship. From this switch, we deduced a combinatorial code including CSPs that mediate both attraction between synaptic partners and repulsion between non-partners^5,6^. The anatomical switch changed the odor response of the new PN partner and markedly increased male-male courtship. We generalized three manipulation strategies from this rewiring— increasing repulsion with the old partner, decreasing repulsion with the new partner, and matching attraction with the new partner—to successfully rewire a second ORN type to multiple distinct PN types. This work demonstrates that manipulating a small set of CSPs is sufficient to respecify synaptic connections, paving ways to explore how neural systems evolve through changes of circuit connectivity.

## Main

The precise wiring of neural circuits is the foundation of brain function. In his chemoaffinity hypothesis, Sperry speculated that “brain cells and fibers must carry some kind of individual identification tags, by which they are distinguished one from another almost, in many regions, to the level of a single neuron”^7^. Many CSPs have since been identified that guide axons to specific target regions^1,2^. CSPs that instruct synaptic partner selection within a specific target region have also begun to be identified^8^. However, disrupting individual CSPs, even with complete loss-of-function mutations, usually leads to partial phenotypes at specific wiring steps, particularly in synaptic partner selection^5,6,9^, suggesting considerable redundancy. Although redundancy could, in principle, increase the robustness of circuit wiring^3^, it poses technical challenges to using a reductionist approach to achieve a complete understanding of how different CSPs work together to assemble a functional circuit, a central goal of developmental neurobiology.

An alternative approach to understanding circuit assembly is to re-engineer the combinatorial expression of CSPs in a single neuron type, with the aim of completely rewiring these neurons away from their endogenous synaptic partner and to a new partner. A challenge of rewiring a neural circuit is that the number of CSPs needed is suspected to be large in general^8^. Here, we report taking such an approach in the *Drosophila* olfactory circuit. In adult *Drosophila*, about 50 types of ORNs form one-to-one synaptic connection with 50 types of PNs at 50 discrete glomeruli, providing an excellent system for studying mechanisms underlying synaptic partner matching. Several recent studies have motivated our attempts to rewire the fly olfactory circuits. First, despite the 3-dimensional organization of 50 glomeruli in adults, during development, each ORN axon only needs to search for synaptic partners along a 1D trajectory on the surface of the antennal lobe^10^. This greatly reduces the number of synaptic partners from which individual ORN axons need to distinguish. Second, examining ORN axon development at single-neuron resolution revealed that each ORN axon extends multiple transient branches along its trajectory in early stages of development and branches that contact partner dendrites are selectively stabilized^4^. Third, in a companion manuscript^5^, we report the identification of three CSP pairs that signal repulsion during the partner matching process to prevent synaptic connections between non-cognate ORN and PN pairs. These repulsive CSPs, along with several attractive CSPs previously characterized^6^ and reported here, are key components in the combinatorial codes for synaptic partner matching we are about to describe.

### Genetic tools to visualize rewiring

We first sought to rewire ORNs that normally target their axons to the DA1 glomerulus (DA1-ORNs) to instead synapse with VA1v-PNs, whose dendrites tile the VA1v glomerulus (Fig. 1a), by combinatorially manipulating the expression levels of different CSPs in DA1-ORNs. We chose these two glomeruli because the axons of both DA1-ORNs and VA1v-ORNs take similar trajectory during development^11^ and because they process signals that have the opposite effects on male courtship activity^12,13^ (see details below). To simultaneously manipulate the expression levels of multiple CSPs only in DA1-ORNs during the wiring process, we generated a genetic driver that specifically labels DA1-ORNs across developmental stages using split-GAL4^14^ (hereafter DA1-ORN driver, Extended Data Fig. 1). To examine the matching of DA1-ORN axons with the dendrites of either DA1-PNs or VA1v-PNs in adults, we co-labeled DA1-ORNs (using the split-GAL4 above) with either DA1-PNs or VA1v-PNs in the same adult brain using the orthogonal QF/QUAS^15^ and LexA/LexAop^16^ systems, respectively (Fig. 1b, c). In wild-type flies, DA1-ORN axons overlapped with DA1-PN dendrites but not with VA1v-PN dendrites (Figs. 1b, c, 2a).

**Fig. 1.**
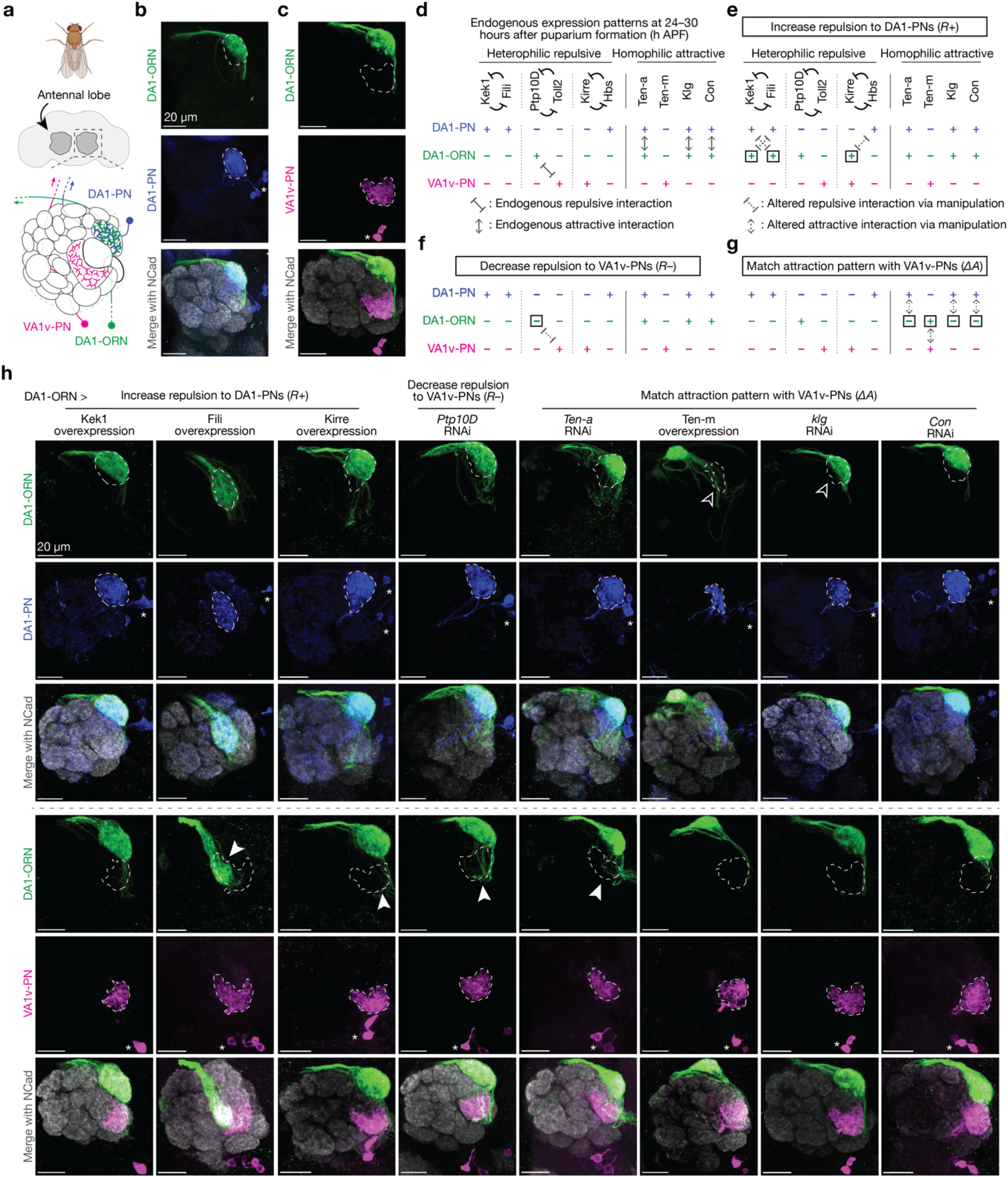
Manipulating single CSPs in DA1-ORNs produces minor DA1-ORNs®VA1v-PNs rewiring. **a**, Adult *Drosophila* brain and antennal lobe schematics. DA1-ORN axons (green) match with DA1-PN dendrites (blue), but not with VA1v-PN dendrites (magenta). Same color code in all other panels. **b**, Maximum z-projection of adult antennal lobes around DA1-ORN axons (green, labeled by a membrane-targeted GFP driven by a split-GAL4) and DA1-PN dendrites (blue, labeled by a membrane-targeted RFP driven by a QF2 driver). DA1 glomerular border (dash-outlined) determined by N-cadherin (Ncad) staining. * designates PN cell bodies; scale bar = 20 µm (all panels). **c**, Same as **b**, but with VA1v-PN dendrites (magenta) labeled instead of DA1-PN dendrites. VA1v glomerular border dash-outlined. **d**, Summary of expression levels of the ten CSPs considered in the rewiring experiments. ‘+’ and ‘–’ indicate relatively high and low expression levels, respectively. The expression levels are largely inferred from the previously collected single-cell RNA sequencing dataset, and are confirmed or corrected with the protein data when available (Extended Data Fig. 3). 24–30h APF is a developmental stage just prior to the onset of synaptic partner selection via stabilization of transient ORN axon branches^4^. **e**, Proposed genetic manipulations (in DA1-ORNs only) to increase the repulsion between DA1-ORN axons and DA1-PN dendrites during development. Square boxes in **e–g** indicate proposed genetic manipulations, with ‘+’ for overexpression and ‘–’ for RNAi knockdown. **f**, Same as **e**, but on proposed genetic manipulations to decrease the repulsion between DA1-ORN axons and VA1v-PN dendrites. **g**, Same as **e**, but on proposed genetic manipulations to match the attraction between DA1-ORN axons and VA1v-PN dendrites. **h**, Rewiring effects when CSPs are manipulated individually. Genetic manipulations are labeled on the top. Maximum z-projections of adult antennal lobes around DA1-ORN axons are shown. Top three rows: DA1-PNs are co-labeled with borders outlined. The open arrowhead indicates the decrease of overlap between DA1-ORN axons and DA1-PN dendrites. Bottom three rows (different brains from the top three rows): VA1v-PNs are co-labeled with borders outlined. Arrowheads indicate the mismatch of DA1-ORN axons with VA1v-PN dendrites. Overlapping ratios are quantified in Fig. 2a.

**Fig. 2.**
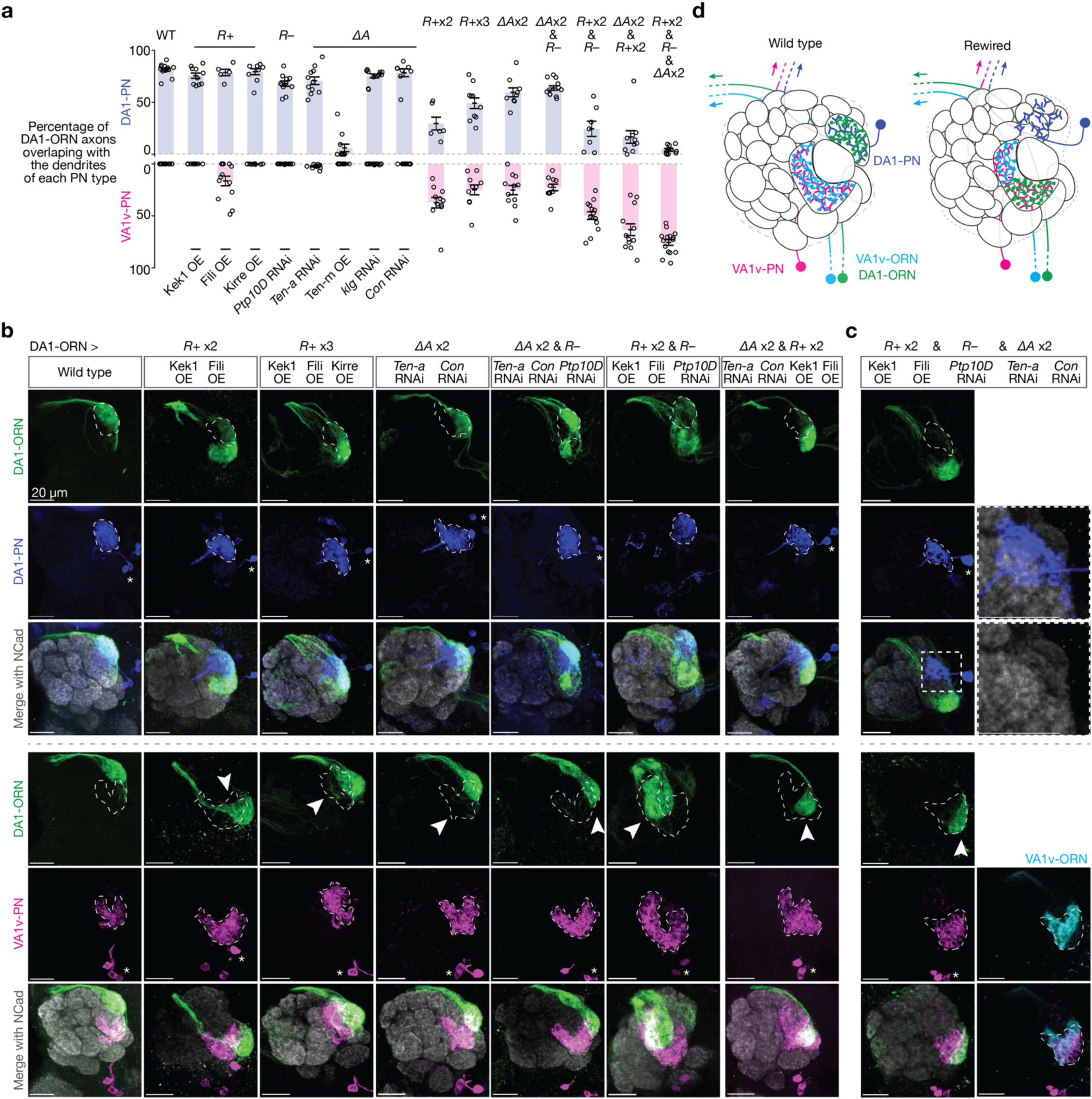
Simultaneous altering the expression of five CSPs in DA1-ORNs causes nearly complete DA1-ORNs®VA1v-PNs rewiring. **a**, Percentage of DA1-ORN axons overlapping with the dendrites of DA1-PNs (top) and VA1v-PNs (bottom). Circles indicate individual antennal lobes; bars indicate the population mean ± s.e.m. ‘*R+* x2’: Kek1 OE + Fili OE. ‘*R+* x3’: Kek1 OE + Fili OE + Kirre OE. ‘*ΔA* x2’: *ten-a* RNAi + *con* RNAi. Same labels used in **b**. **b**, Rewiring effects when CSPs are manipulated combinatorially. Genetic manipulations are labeled on the top. Maximum z-projections of adult antennal lobes around DA1-ORN axons (green) are shown. Top three rows: DA1-PNs (blue) are co-labeled with borders dash-outlined. Bottom three rows: VA1v-PNs (magenta) are co-labeled with borders dash-outlined. Arrowheads indicate the mismatch of DA1-ORN axons with VA1v-PN dendrites; scale bar = 20 µm; * designates PN cell bodies. Overlapping ratios are quantified in **a**. The leftmost column is a repeat of Fig. 1b, c for ease of comparison within this panel. **c**, Same as **b**, but with all three manipulation strategies combined. Two images on the top of the right column are zoom-ins from the dashed squares to the left. Two images on the bottom of the right column are from the same brain as in the left column, but with VA1v-ORNs co-labeled (cyan, *Or47b* promotor driven rat CD2). **d**, Summary of DA1-ORNs and -PNs as well as VA1v-ORNs and -PNs in the wild-type (left) and DA1-ORN rewired (right) antennal lobe. In the rewired lobe: axons of DA1-ORNs and VA1v-ORNs split VA1v-PN dendrites; DA1-PN dendrites spread into multiple adjacent glomeruli.

### Three manipulation strategies for rewiring

To achieve rewiring, we considered 10 CSPs that are likely to signal attractive or repulsive interactions during ORN-PN synaptic partner matching (Fig. 1d). Ten-a and Ten-m are type II transmembrane proteins that each exhibits matching expression patterns across ORN and PN types and mediate homophilic adhesion^6^. Connectin (Con) and Klingon (Klg) are also homophilic adhesion molecules involved in the development of *Drosophila* visual and neuromuscular circuit, respectively^17,18^. Based on single-cell RNA sequencing (scRNAseq) data^19,20^, we found that Con and Klg also exhibited matching expression patterns across ORN and PN types (Extended Data Fig. 2). RNAi-knocking down^21,22^ of *con* and overexpression of Klg caused partial mismatching phenotypes consistent with their promoting homophilic attraction between ORNs and PNs (Extended Data Fig. 2). The remaining 6 CSPs form three groups—Kekkon 1(Kek1) with Fish-lips (Fili), Protein tyrosine phosphatase 10D (Ptp10D) with Toll2, and Kin of irre (Kirre) with Hibris (Hbs)—and signal repulsion between ORNs and PNs^5,9^.

The expression level of all CSPs were largely inferred from scRNAseq data sets during development^19,20^. Since the scRNAseq data are prone to measurement noise and may not accurately reflect protein expression due to post-transcriptional regulation, we corrected our RNA data using protein data and *in vivo* genetic manipulation results in CSPs where additional data were available (Fig. 1d, Extended Data Fig. 3). As summarized in Fig. 1d, developing DA1-ORNs and DA1-PNs in wild type contained attractive interactions from three CSPs (Ten-a, Klg, and Con) but no repulsive interaction from the three repulsive pairs, in accordance with them forming synaptic partners in adults (Fig. 1d, Fig. 2a). By contrast, developing DA1-ORNs and VA1v-PNs contained no attractive interactions from the four attractive CSPs but repulsive interactions from one CSP pair (Ptp10D and Toll2) (Fig. 1b, Extended Data Fig. 3), consistent with them being non-synaptic partners in adults (Fig. 1c, Fig. 2a).

To facilitate rewiring, we utilized three strategies of genetic manipulation during development, all restricted only to DA1-ORNs (Fig. 1e–g). (1) We increased the repulsion strength between DA1-ORN axons and DA1-PN dendrites (‘*R+*’) to destabilize their interaction. Since repulsive CSPs Kek1, Fili, and Hbs are highly expressed in wild-type DA1-PNs, we overexpressed their interaction partners Fili, Kek1, and Kirre in DA1-ORNs (Fig. 1e). (2) We decreased the strength of repulsion between DA1-ORN axons and VA1v-PN dendrites (‘*R–*’) to stabilize their interaction. Since Ptp10D from DA1-ORNs mediates the repulsive interaction with Toll2 from VA1v-PNs in wild-type flies, we knocked down *Ptp10D* expression in DA1-ORNs (Fig. 1f). (3) We matched the expression pattern of attractive molecules between DA1-ORN axons and VA1v-PN dendrites (‘*ΔA’*) to stabilize their interactions and at the same time to destabilize the interactions between DA1-ORNs and DA1-PNs. Since the expression pattern of none of the four attractive CSPs between DA1-ORNs and VA1v-PNs match in wild-type flies, we genetically manipulated all four of them independently (Fig. 1g).

### Single-CSP manipulations cause minor rewiring

To start, we used the DA1-ORN driver (Extended Data Fig. 1) to overexpress or knockdown different CSPs in DA1-ORNs and to examine their individual effects on synaptic partner matching. All transgenes used in the repulsive interactions were validated in the companion study^5^. All transgenes used in the attractive interactions were either used in previous studies^6,17,18^ or confirmed via multiple RNAi lines (Extended Data Fig. 2). Across the eight single-CSP manipulations (Fig. 1e–g), six showed observable but subtle DA1-ORNs→VA1v-PNs mismatching phenotypes (middle six columns in Fig. 1h, quantified in Fig. 2a and Extended Data Fig. 4), consistent with the results from the previous manipulation experiments using these CSPs^4–6,9^. In the ‘Ten-m overexpression’ manipulation, most DA1-ORN axons no longer overlapped with DA1-PN dendrites, but none of the mistargeted DA1-ORN axons overlapped with VA1v-PN dendrites. This is consistent with the previous finding that Ten-m-overexpressing DA1-ORN axons most likely mismatch with DL3-PN dendrites^4^. This could be because the CSP profile of DL3-PNs matches with the profile of DA1-ORNs (after Ten-m overexpression) better than that of DA1-PNs.

### Combinatorial manipulations enhance rewiring

Next, we simultaneously manipulated the expression of multiple CSPs in DA1-ORNs. We first aimed to find the CSP combination within each of the three manipulation strategies described above (*R+*, *R–*, and *ΔA*; Fig. 1e–g) that can most strongly decrease the overlap between DA1-ORN axons and DA1-PN dendrites (loss of innervation, LoI) and increase the overlap between DA1-ORN axons and VA1v-PN dendrites (gain of innervation, GoI). Overexpression of both Kek1 and Fili (‘*R+* x2’ in Fig. 2a, b) led to a significant LoI and a significant GoI (Extended Data Fig. 4) compared to overexpressing either alone. This was the largest phenotype we observed among the different combinations of overexpressing repulsive CSPs. For example, overexpressing all three repulsive CSPs (Kek1, Fili, and Kirre, ‘*R+* x3’ in Fig. 2a, b) improved neither GoI nor LoI compared to ‘*R+* x2’ (Extended Data Fig. 4). Therefore, we chose overexpressing Kek1 and Fili (‘*R+* x2’) as the best combination for the strategy of increasing repulsion between DA1-ORNs and DA1-PNs. Similarly, for the strategy of matching the expression pattern of attractive molecules between DA1-ORNs and VA1v-PNs, we found that knocking down *Ten-a* and *Con* simultaneously yielded the most significant LoI and GoI (‘*ΔA* x2’ in Fig. 2a, b). We chose knocking down *Ptp10D* (‘*R–*’ in Fig. 2a, b) as the strategy of decreasing the repulsion of DA1-ORNs and VA1v-PNs since it is the only manipulation available here.

Next, we combined the best options from the three manipulation strategies. When we used two strategies simultaneously (‘*ΔA* x2 & *R–*’, ‘*ΔA* x2 & *R+* x2’, and ‘*R+* x2 & *R–*’ in Fig. 2a, b), in most cases, the LoI further decreased and the GoI further increased compared to each strategy alone. For example, in ‘*ΔA* x2 & *R+* x2’, the LoI was significantly more severe than the LoI in either ‘*ΔA* x2’ or ‘*R+* x2’, and the GoI in the combined group was also significantly larger than the GoI from each group (Fig. 2a, b, Extended Data Fig. 4).

When we combined all three manipulation strategies, nearly all DA1-ORN axons disconnected with DA1-PN dendrites and overlapped with VA1v-PN dendrites (‘*R+* x2 & *R–* & *ΔA* x2’ in Fig. 2a, c, d). Dendrites of DA1-PNs appeared to spread into multiple adjacent glomeruli (inset in Fig. 2c), potentially forming synaptic connections with new ORN partners^10^. Further, DA1-ORN axons only overlapped with part of VA1v-PN dendrites (bottom of Fig. 2c). We confirmed that the non-overlapping part of VA1v-PN dendrites matched with their natural partner VA1v-ORN axons (bottom of Fig. 2c), presumably because we did not genetically manipulate either VA1v-ORNs or VA1v-PNs. Notably, the axons of DA1-ORNs and VA1v-ORNs are segregated in the rewired flies (Fig. 2c), suggesting potential axon-axon repulsive interactions as previously shown in a different context^23^.

In this final rewiring experiment (referred hereafter as DA1-ORN rewired flies), the expression levels of five CSPs were changed in DA1-ORNs (Kek1, Fili, Ptp10D, Ten-a, and Con; Fig. 2c). When any one of the five CSP changes was omitted, the rewiring was less complete (Extended Data Fig. 5). Despite the fly DA1 glomerulus being sexually dimorphic in size^24,25^, the rewiring of DA1-ORN axons to VA1v-PN dendrites showed similar levels of change in male and female flies (Extended Data Fig. 6). Moreover, axons of several additional types of ORNs remained confined within their original glomeruli in rewired flies (Extended Data Fig. 6), supporting that the rewiring is specific to the DA1 and VA1v glomeruli.

### Rewiring alters VA1v-PN odor response

To examine whether the anatomical rewiring of DA1-ORN axons to VA1v-PN dendrites is accompanied with the formation of functional synaptic connections, we measured the neural response of VA1v-PN dendrites to VA1v- or DA1-specific odors in tethered flies (Fig. 3a). All ORN-PN connections are excitatory and use the same cholinergic neurotransmitter system^26^. We used the LexA/LexAop system to express GCaMP7b in VA1v-PNs and measured intracellular Ca^2+^ concentrations via two-photon excitation of GCaMP7b^27^ as a proxy for neural activity. We simultaneously expressed and co-imaged tdTomato in DA1-ORNs with GCaMP7b and confirmed the occurrence of DA1-ORNs→VA1v-PNs rewiring in these flies (Fig. 3a).

**Fig. 3.**
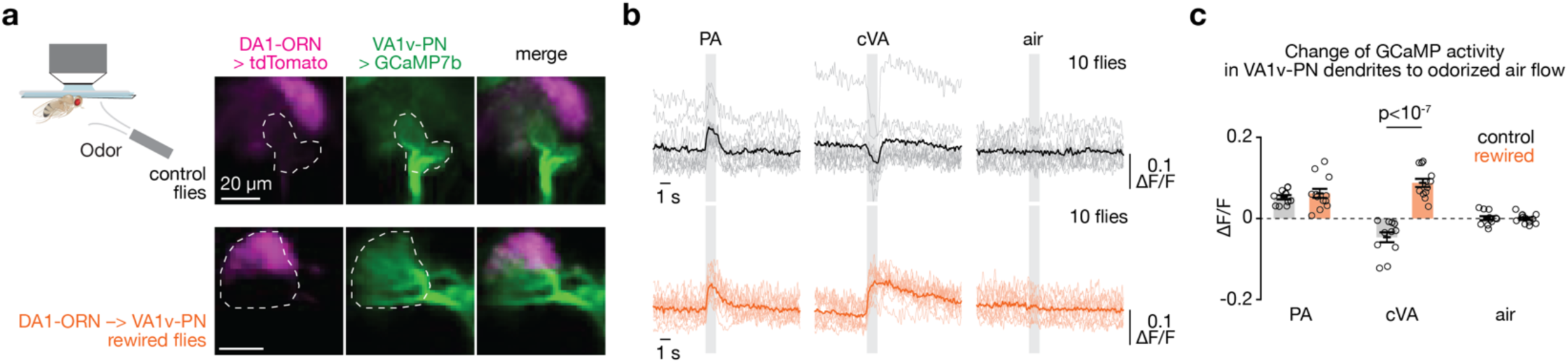
VA1v-PNs in DA1-ORN rewired flies respond to both VA1v- and DA1-specific odors. **a**, Imaging neural activity in a plate-tethered fly with odorized air flow delivered to the fly antennae. Images of tdTomato signal in DA1-ORN axons and GCaMP7b signal in VA1v-PN dendrites are shown from a control fly (top) and a DA1-ORN rewired fly (bottom). Images are averaged across the entire recording. The VA1v glomerulus is outlined according to GCaMP7b signal. Scale bar = 20 µm. Note that the imaging angle here is from dorsal to ventral whereas all the other staining images are from anterior to posterior. **b**, Averaged GCaMP7b activity in VA1v-PN dendrites in response to odorized air flows, measured by fluorescence intensity change over baseline (ΔF/F). Top: control flies. Bottom: DA1-ORN rewired flies. The grey vertical stripes indicate odorized air flows (1 s each). Light-colored traces indicate the means of individual flies; dark-colored traces indicate the population mean. In wild-type flies, palmitoleic acid (PA) specifically activates VA1v-ORNs^12^ and 11-*cis*-vaccenyl acetate (cVA) specifically activates DA1-ORNs^13,28,29^. **c**, Change of GCaMP7b activity in VA1v-PN dendrites to odorized air flows. The change of activity is calculated by subtracting the average GCaMP7b activity in the 0.5 s before the onset of odor delivery from that in the last 0.5 s of odorized airflow. Circles indicate the means of individual flies; bars indicate the population mean ± s.e.m. Unpaired t-test is used.

We next tested odor responses of VA1v-PN dendrites. 11-*cis*-vaccenyl acetate (cVA) is a pheromone that specifically activates DA1-ORNs in the fly antennal lobe^13,28,29^. Palmitoleic acid (PA) is a fly cuticular pheromone that specifically activates VA1v-ORNs in the fly antennal lobe^12^. In wild-type flies, we found that the dendrites of VA1v-PNs increased response to PA and decreased response to cVA (Fig. 3b, c). The inhibitory response of VA1v-PNs to cVA in wild-type flies is consistent with the previously described lateral inhibition from local interneurons in the fly olfactory circuit^30,31^. In the rewired flies, however, both PA and cVA activated VA1v-PNs (Fig. 3b, c), supporting functional synaptic connections between DA1-ORN axons and VA1v-PN dendrites. We cannot rule out the possibility that altered connectivity of local interneurons, which exhibit diverse anatomical patterns^32,33^, also contributes to the altered odor response. However, the inhibitory response of VA1v-PNs to odors that do not strongly activate VA1v- or DA1-ORNs remained similar between the rewired flies and wild-type flies (Extended Data Fig. 7), supporting that the connection between VA1v-PNs and local interneurons remained largely unchanged.

### Rewiring promotes male-male courtship

Would DA1-ORNs→VA1v-PNs rewiring lead to any behavioral change in flies? In *Drosophila melanogaster*, cVA is only produced in males and acts through the Or67d odorant receptor expressed in DA1-ORNs to inhibit the courtship of males towards other males or recently mated females^13^ (due to cVA transferred from males to females during copulation^34^). The pheromone PA, on the other hand, promotes courtship in males through the Or47b odorant receptor expressed in VA1v-ORNs^12^. Therefore, in rewired flies, a pheromone that normally inhibits male-male courtship (cVA) now activates a pathway (VA1v) that promotes courtship. This predicts that rewired males may attempt to court other males.

To test this prediction, we introduced two virgin males—one wild type and one with DA1-ORN rewired— into the same behavioral chamber (Fig. 4a). We then recorded video for 25 minutes and analyzed from both males the unilateral-wing-extension events (Fig. 4b, Supplementary Videos 1, 2), a typical male courtship behavior during which males vibrate one of their wings to produce courtship song^35^. We found that the rewired males exhibited unilateral wing extensions towards their wild-type partner males significantly more frequently than the other way around (Fig. 4c, d). In a separate experiment, we introduced one male—either wild type or rewired— with a virgin female into the behavioral chamber. We did not observe any detectable differences in the courtship activity towards virgin females in wild-type and rewired males (Extended Data Fig. 8a-c). This is consistent with our working model since a virgin female does not have cVA, and connections between VA1v-ORNs and VA1v-PNs in rewired flies remained intact as assayed anatomically (Fig. 2c) and physiologically (Fig. 3b, c). Experimental silencing or activation of DA1-ORNs in rewired males further revealed that both the loss-of-connection to DA1-PNs and the gain-of-connection to VA1v-PNs in rewired males contributed to the increased male-male courtship activity (Extended Data Fig. 8d–l). Finally, when five virgin rewired males were introduced into the same behavioral chamber, they exhibited vigorous chasing and courtship activities, sometimes forming a courtship chain in which a male attempted to court the male in front of him while being courted by another male behind him (Supplementary Video 3).

**Fig. 4.**
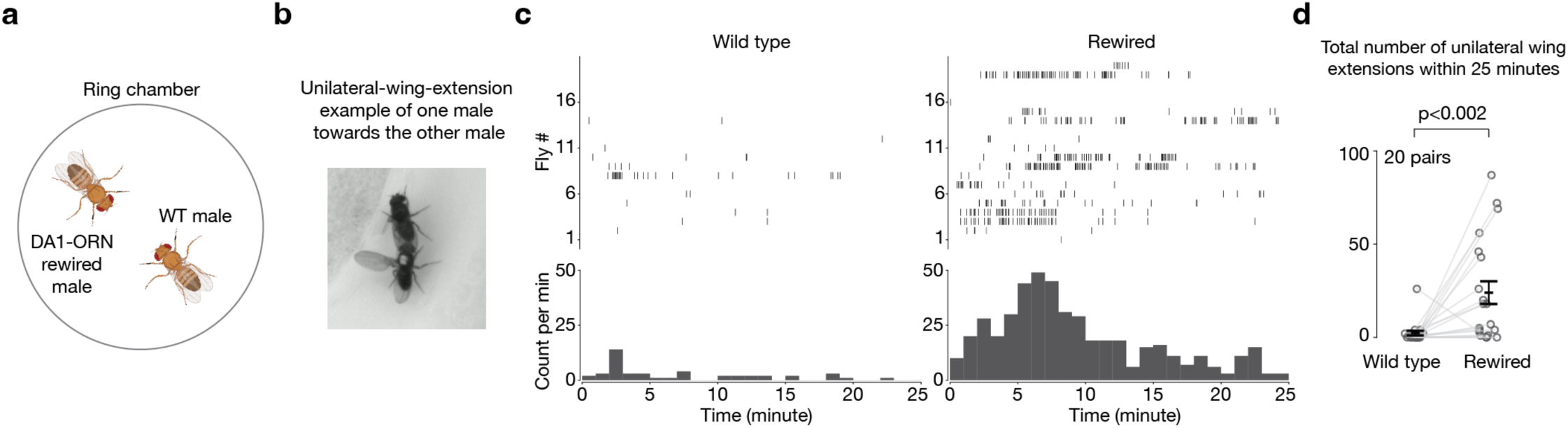
Male flies with DA1-ORNs rewired show elevated courtship activity towards other males. **a**, Courtship assay where one wild-type male and one DA1-ORN rewired male are introduced in the same behavioral chamber to monitor their courtship activity towards each other. The chamber diameter is 2 cm. **b**, Example frame of unilateral wing extension from a DA1-ORN rewired male (white dot on the thorax) towards a wild-type male. **c**, Rasters of unilateral wing extensions (top) and extension count per minute (1-min bins, bottom). Left: wild-type males; right: DA1-ORN rewired males. The same fly number in the left and right panels are the fly pair from the same experiment. **d**, Total unilateral-wing-extension number within 25-minute recordings. Circles indicate individual flies; bars indicate the population mean ± s.e.m. Wilcoxon signed-rank test is used given the non-normal distribution of the data points.

### Generalization to other glomeruli

Do the same set of CSPs and wiring strategies also apply to the ORN-PN synaptic partner matching in other glomeruli? We tested this by attempting to rewire the axons of another ORN type, VA1d-ORNs, to the dendrites of PNs targeting three distinct neighboring glomeruli: VA1v, DC3, and DL3 (Fig. 5).

**Fig. 5.**
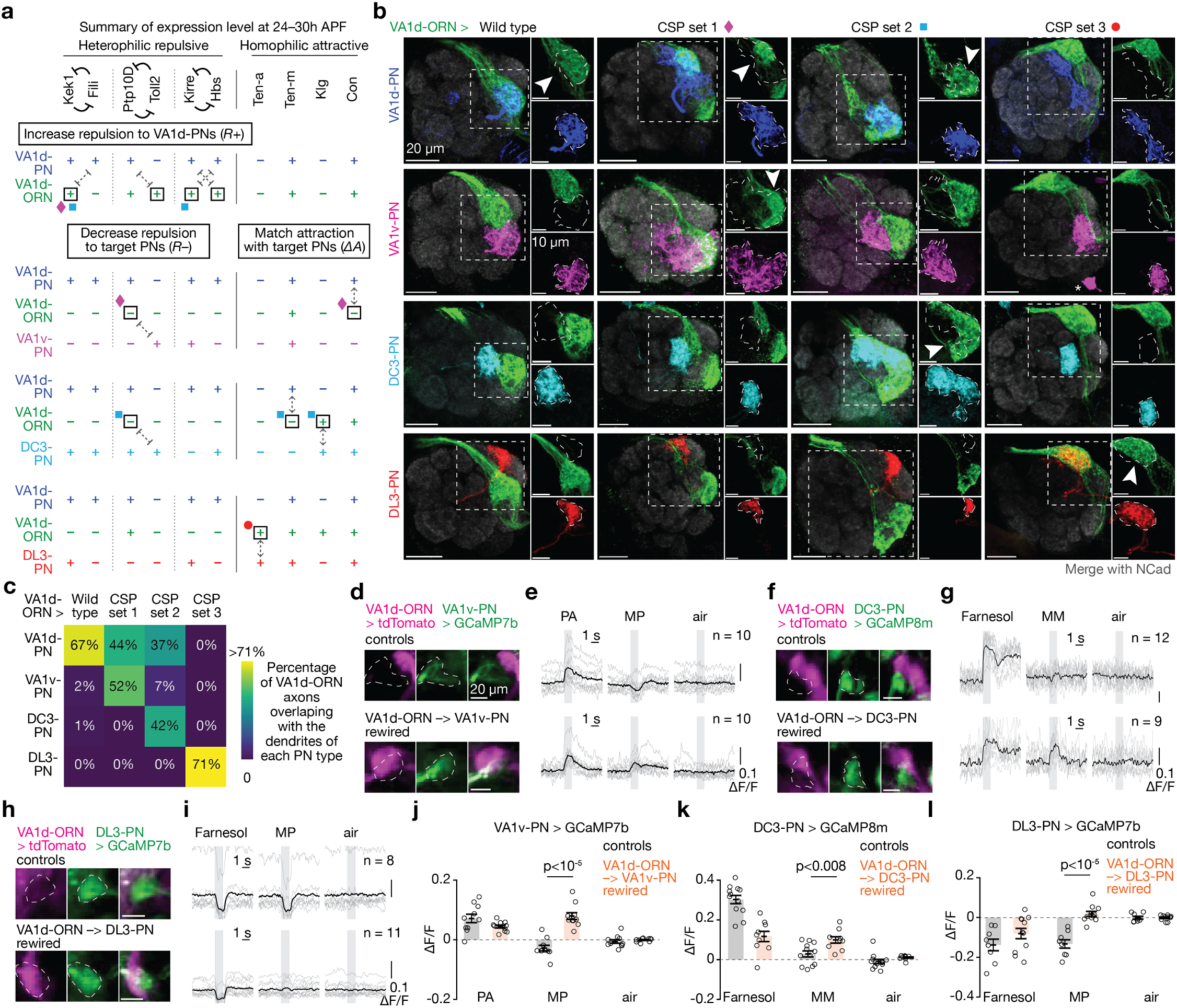
Rewiring VA1d-ORNs with distinct PN partners following the same manipulation strategies. **a**, Summary of expression levels of the ten CSPs and the three manipulation strategies considered in the experiments of rewiring VA1d-ORNs (green) away from VA1d-PNs (blue) and towards VA1v-PNs (magenta), DC3-PNs (cyan), or DL3-PNs (red), respectively. Same color code here and in (**b**). Nomenclature same as Fig. 1d–g. Available protein expression data are in Extended Data Fig. 9. Diamonds, squares, and circles indicate the genetic manipulations used in the final switches of VA1d-ORNs to VA1v-, DC3-, and DL3-PNs, respectively. See main text for more details. **b**, Maximum z-projections of adult antennal lobes around VA1d-ORN axons in wild type and all the three rewired experiments. In each row, VA1d-ORNs are co-labeled with one PN type according to labels on the left. In each column, the same set of genetic manipulations is used. CSP set 1 = Kek1 OE + *Ptp10D* RNAi + *con* RNAi + *sema-2b* RNAi; CSP set 2 = Kek1 OE + Kirre OE + *Ptp10D* RNAi + *Ten-m* RNAi + Klg OE; CSP set 3 = Ten-a OE + *sema-2b* RNAi. *sema-2b* RNAi is used for rerouting the VA1d-ORN trajectory to better match the dendrites of VA1v-PNs and DL3-PNs. See main text and Extended Data Fig. 10 for detail. Borders of the dendrites of each PN type are dash-outlined. Arrowheads indicate the overlap of VA1d-ORN axons with the dendrites of specific PN types. Insets are zoom-ins from the dashed squares to their left. Scale bars are 20 µm in large images and 10 µm for insets. **c**, Percentage of VA1d-ORN axons overlapping with the dendrites of each PN type (indicated on the left) in wild type and the three rewired conditions (indicated on the top). n ≥ 6 for all the conditions. See Extended Data Fig. 10 for detail. **d**, Images of tdTomato signal in VA1d-ORN axons and GCaMP7b signal in VA1v-PN dendrites from a control fly (top) and a VA1d-ORN®VA1v-PN fly (bottom). Images are averaged across the entire recording. The VA1v glomerulus is outlined according to GCaMP7b signal. Scale bar = 20 µm. Same imaging setup as in Fig. 3. **e**, Averaged GCaMP7b activity in VA1v-PN dendrites in response to odorized air flows. Top: control flies. Bottom: VA1d-ORN®VA1v-PN flies. The grey squares indicate odor deliveries (1s each). Grey lines indicate the means of individual flies; black lines indicate the population mean. In wild-type flies, fly pheromone palmitoleic acid (PA) specifically activates VA1v-ORNs^12^ and fly pheromone methyl palmitate (MP) mainly activates VA1d-ORNs^28^. **f, g,** Same as **d, e**, but examining rewiring of VA1d-ORNs to DC3-PNs instead of VA1v-PNs. GCaMP8m is used instead of GCaMP7b. In control flies, odorant farnesol mainly activates DC3-ORNs^36^. Fly pheromone methyl myristate (MM) specifically activate VA1d-ORNs and VA1v-ORNs^12^ and is used here instead of MP since we observed positive response of DC3-PNs to MP in wild type (unpublished observation). **h, i**, Same as **d, e**, but examining rewiring of VA1d-ORNs to DL3-PNs instead of VA1v-PNs. Note that farnesol is used as a control since DL3-specific odorants remain unknown. In the control fly image, VA1d-ORN signal is absent because VA1d-ORN axons and DL3-PN dendrites occupy different z positions from the current imaging perspective. **j–l**, Change of GCaMP activity in dendrites of VA1v-PNs (**j**), DC3-PNs (**k**), and DL3-PNs (**l**) to odorized air flow. The change of activity is calculated by subtracting the average GCaMP activity in the 0.5 s before the onset of odor delivery from the average GCaMP activity in the last 0.5 s of odorized air flow. Circles indicate the means of individual flies; bars indicate the population mean ± s.e.m. Same number of flies as shown in **e**, **g**, and **i**, respectively. Unpaired t-test is used.

We used a genetic driver that specifically labels VA1d-ORNs across developmental stages using split-GAL4^4^, and simultaneously labeled the dendrites of either VA1d-PNs, VA1v-PNs, DC3-PNs, or DL3-PNs in the same adult brain using the orthogonal LexA/LexAop^16^ or QF/QUAS^15^ systems (Fig. 5b). In wild-type flies, VA1d-ORN axons almost exclusively overlapped with the dendrites of VA1d-PNs and showed minimal overlap with the dendrites of other PN types (Fig. 5b, c). The goal of rewiring is to switch the axons of VA1d-ORNs to match with the dendrites of each of the three other PN types in individual experiments.

Based on the same 10 CSPs described above, during the development of wild-type flies, VA1d-ORN axons and VA1d-PN dendrites form two attractive interactions (via Ten-m and Con) and no repulsive interactions (Fig. 5a, Extended Data Fig. 9). For the first manipulation strategy that aims to increase the strength of repulsion between VA1d-ORNs and VA1d-PNs, we could overexpress repulsive CSPs Kek1, Toll2, Kirre, or Hbs in all three rewiring attempts (Fig. 5a, top). For the second manipulation strategy that aims to decrease the strength of repulsion between VA1d-ORNs and other PN types, we sought to knock down *Ptp10D* in two of the three switch attempts and do nothing in the switch attempt to DL3-PNs since DL3-PNs do not exhibit any repulsive interactions with VA1d-ORNs from these three repulsive pairs (Fig. 5a, bottom left). For the third manipulation strategy that aims to match the expression pattern of attractive molecules between VA1d-ORNs and other PN types, we could overexpress or knock down the expression of these four attractive CSPs accordingly (Fig. 5a, bottom right).

By combining the different combinations of manipulations described above, we were able to rewire more than half of VA1d-ORN axons to match with the dendrites of either VA1v-PNs or DC3-PNs in two separate experiments, and to rewire almost all VA1d-ORN axons to match with DL3-PN dendrites in a third experiment (Fig. 5b, c). In all three rewiring experiments, the part of VA1d-ORN axons that did not match with the dendrites of target PNs remained matching with dendrites of their natural partner VA1d-PNs (Fig. 5b, c). Note that in the rewiring to VA1v-PNs and DL3-PNs, we also included an additional manipulation, Sema-2b knockdown. This is because VA1d-ORNs have a higher expression level of *sema-2b* than VA1v-ORNs and DL3-ORNs^19^. Based on this, we speculated that VA1d-ORN axons take a more dorsolateral trajectory than VA1d-ORNs when they sweep through the antennal lobe surface. Since a single ORN axon mainly searches in the vicinity of their trajectory^4^, we included *Sema-2b* knockdown to shift the axons of VA1d-ORNs more dorsolaterally^10,11^ so their trajectories could be closer to the dendrites of VA1v-PNs and DL3-PNs. Consistently, when all the manipulations remained the same but leaving out the *Sema-2b* knockdown, there was less matching between VA1d-ORN axons and VA1v- or DL3-PN dendrites (Extended Data Fig. 10).

To test whether the anatomical rewiring of VA1d-ORNs described above lead to the formation of functional synaptic connections, we examined in rewired flies if the different PN types would gain responses to VA1d-ORN-specific odors, pheromones methyl palmitate (MP)^28^ or methyl myristate^12^ (see Fig. 5 legend for more detail). Using the same setup as in Fig. 3a, we measured the neural response of VA1v-, DC3-, and DL3-PNs, separately, via two-photon excitation of GCaMP variants expressed using the LexA/LexAop or QF/QUAS system in these PNs (Fig. 5d–i). We also co-expressed tdTomato in VA1d-ORNs to confirm the anatomical switch in these flies. In all three rewiring experiments, the dendrites of target PNs gained response to VA1d-ORN-specific odors compared to in wild-type flies (Fig. 5d–l). Note that in the case of DL3 (Fig. 5i, l), although rewiring eliminated the inhibitory response of DL3-PNs to MP, the magnitude of the positive response was much smaller. We speculate that this could result from substantial lateral inhibition that DL3-PNs might receive from other MP-responding ORN types. Altogether, these results demonstrate that the three genetic strategies for altering the cell-surface combinatorial code are generalizable for selecting synaptic partners in the fly olfactory circuit.

## Discussion

Here, we demonstrated that the fly olfactory circuit could be largely rewired when two to five cell-surface proteins (CSPs) were changed in a single ORN type (Figs. 1, 2, and 5). This occurred even though dozens of CSPs are differentially expressed between different ORN types during the synaptic partner matching period (Extended Data Fig. 3). The rewiring expanded the physiological response to odors in downstream PNs (Figs. 3 and 5) and altered the courtship behavior in one case (Fig. 4).

The CSP combinatorial code for rewiring should be closely related, if not identical, to the CSP code used during natural wiring. To illustrate, consider the rewiring of DA1-ORNs®VA1v-PNs. First, the five CSPs involved in the rewiring are differentially expressed between DA1-ORNs and VA1v-ORNs (Extended Data Fig. 3). The directions of gene expression manipulation—whether up- or down-regulation—match the discrepancy between these two ORN types. Second, both loss- and gain-of-function manipulations in most of the five CSPs alone significantly decreased the matching of DA1-ORN axons with DA1-PN dendrites or caused mismatch of DA1-ORN axons with VA1v-PN dendrites (Extended Data Fig. 4), supporting the involvement of these CSPs in distinguishing the wiring specificity of DA1-ORNs and VA1v-ORNs naturally. Finally, rewiring leads to a gain-of-function at both the physiological and behavioral levels, suggesting its potential utility in an evolutionary context.

The results that the rewiring could be successful despite our lack of precise control over the level and timing of the CSP manipulations suggests that the combination of key CSPs is more critical than the exact levels and timing of their expression. This is consistent with the general notion that many biological systems are robust in their tolerance to variations in gene expression. The precision of rewiring may be further improved if we could better control our genetic manipulations in level and timing, and by manipulating additional CSPs that we may have missed (e.g., in the case of VA1d-ORNs®DA1-PNs and VA1d-ORNs®DC3-PNs in Fig. 5).

Our results show that synaptic partner matching appears flexible in the specific CSPs used so long as they execute a common set of strategies: matching attractive CSPs between partners, avoiding repulsive CSPs between partners, and displaying repulsive CSPs between non-partners. Furthermore, CSPs of different families^5,6,17,18^— those containing immunoglobulin-like domains (Klg, Kirre, Hbs, Kek1), leucine-rich repeats (Con, Fili, Kek1, Toll2), fibronectin III domains (Ptp10D, Hbs), and teneurin (Ten-a, Ten-m)—work together in different combinations for synaptic partner matching at different glomeruli. We speculate that these different protein families may converge onto common intracellular signaling pathways to regulate cytoskeletal changes that underlie attraction^4^ and repulsion. We further note that protein motifs in these CSPs, and in many cases individual CSPs themselves, are evolutionarily conserved across invertebrates and vertebrates^2,8^. Thus, the combinatorial action of different CSP types we described here may be used to control synaptic partner matching in nervous systems from insects to mammals.

## Methods

### Fly husbandry and stocks

Flies were reared on a standard cornmeal medium at 25°C under a 12-hour light and 12-hour dark cycle. To enhance transgene expression levels, flies from all genetic perturbation experiments, including control groups, were shifted to 29°C shortly before puparium formation. Detailed genotypes for each experiment are listed in Extended Data Table 1.

### Molecular cloning and generation of transgenic flies

To generate QF2 lines, we used *pENTR/D-TOPO* vectors with different enhancer insertions (gifts from G. Rubin lab) as entry vectors for Gateway cloning into the *pBPQF2Uw* vector using LR Clonase II Enzyme mix (Invitrogen, 11791020). *pBPQF2Uw* was made using NEBuilder HiFi DNA assembly master mix (New England Biolabs) to replace the GAL4 on *pBPGAL4.2Uw-2* vector (Addgene #26227) with QF2 from *pBPGUw-HACK-QF2* (Addgene #80276). The resulting constructs were sequence-verified and inserted into JK22C landing sites by Bestgene. *pGP-5XQUAS-IVS-Syn21-jGCaMP8m-p10* was made using NEBuilder HiFi DNA assembly master mix (New England Biolabs) to replace the 20XUAS on *pGP-20XUAS-IVS-Syn21- jGCaMP8m-p10* vector (Addgene #162387) with 5XQUAS from pQUAST (Addgene #24349). Plasmids were injected to embryos at Bestgene. Genetic labeling with these drivers is unlikely to disrupt normal development, as a previous study showed that drivers with improved translation efficiency could elevate GFP expression by 20 fold with no apparent effect on neuronal morphology^37^.

### Immunostaining

The procedures used for fly dissection, brain fixation, and immunostaining were described previously^10^. For primary antibodies, we used rat anti-DNcad (1:30, from DSHB, RRID # AB_528121), chicken anti-GFP (1:1000, from Aves Labs, RRID # AB_10000240), rabbit anti-DsRed (1:500, from Takara Bio, RRID # AB_10013483), and mouse anti-rat CD2 (1:200; OX-34, Bio-Rad).

### Confocal imaging

Immunostained brains were imaged using a laser-scanning confocal microscope (Zeiss LSM 780). Images of antennal lobes were taken as confocal stacks with 1-mm-thick sections. Representative single sections were shown to illustrate the arborization features of ORN axons and PN dendrites, with brightness adjustment, contrast adjustment, and image cropping done in ImageJ.

### Calculating the percentage of ORN axons matching with PN dendrites

PN dendritic pixels and ORN axonal pixels were defined by first smoothening the image using ‘gaussian blur’ (radius = 2 pixels) and then thresholding the image based on the algorithm ‘Otsu’ in Fiji. We found that this algorithm could efficiently separate the neurons of interest from the background. Irrelevant signals (such as the PN axons, cell bodies, or autofluorescence) that still persisted after the above operations were manually masked out in the analysis. A portion of ORN axons was considered as matching with PN dendrites if they have overlapping pixels on a single *z*-plane in the image. Note that the definition of glomerulus becomes vague as ORN axons and PN dendrites innervate more and more outside the original glomerulus.

The calculated overlap between ORN axons and PN dendrites is always lower than 100%. This is because ORN axons or PN dendrites do not occupy the entire glomerulus because of a technical reason and a biological reason. Technically, if one examines axons and dendrites with super resolution, they should not overlap at all as each physical space should be occupied by only one entity if the resolution is sufficiently high. In our quantifications, we used ‘gaussian blur’ to best recapitulate the adjacent areas of a single axon or dendrites that should be considered as ‘overlap’. This is an empirical parameter and would not achieve 100% overlap. Biologically, besides ORN-PN synapses, both ORNs and PNs also form reciprocal synapses with antennal lobe local interneurons (LNs). Regions with ORN-LN synapses lack PN dendrites; regions with PN-LN synapses lack ORN axons. Thus, ORN axons and PN dendrites don’t overlap in these regions.

In our analyses, we use the same parameters to quantify all genetic conditions. Thus, our conclusions about the changing of ORN-PN overlap under different genetic conditions should not be affected by the above factors.

### Ca^2+^ imaging and data analysis

#### Odor stimuli delivery

10 µL of odorant palmitoleic acid (Thermo Fisher, 376910010) and 11-*cis*-vaccenyl acetate (cVA; Cayman Chemical, 10010101) was applied to filter paper (Amazon, B07M6QJ2JX) inserted inside a 1-ml pipette tip. The pipette tip was left aside for at least half an hour before being positioned approximately 5 mm away from the fly antenna. Close positioning of palmitoleic acid and cVA is necessary because both odorants are large pheromone molecules with relatively low volatility. This method has also been utilized in other studies^12^. Other odorants, such as methyl palmitate (Thermo Fisher, L05509.36), methyl myristate (Thermo Fisher, 165015000), and farnesol (Thermo Fisher, 119121000), were stored in a small glass bottle and delivered to the fly antenna via tubing with a 10% dilution in heavy mineral oil on the day of experiments. A constant stream of charcoal-filtered air (1 L/min) was directed towards the fly, switching to odorant-containing air for 1 second as the odor stimulus before returning to the air stream. A pulse of charcoal-filtered air served as a negative control. Odorants, including the control pulse, were interleaved with at least 15-second intervals. Each odorant was delivered two to three times per recording, with the delivery sequence shuffled within each cycle. As described previously^38^, we glued flies to a custom stage. Dissection and imaging protocols also followed a previous study^38^.

#### Data acquisition and alignment

We used a two-photon microscope with a moveable objective (Ultima IV, Bruker). The two-photon laser (Chameleon Ultra II Ti:Sapphire, Coherent) was tuned to 925 nm in all of the imaging experiments. We used a ×16/0.8 NA objective (Nikon) for all imaging experiments. The laser intensity at the sample was 15–30 mW. A 575-nm dichroic split the emission light. A 490–560-nm bandpass filter (Chroma) was used for the green channel and a 590–650-nm bandpass filter (Chroma) was used for the red channel. We recorded all imaging data using single z plane, at a rate of 9–13 Hz. We perfused the brain with extracellular saline composed of (in mM) 103 NaCl, 3 KCl, 5 N-Tris(hydroxymethyl) methyl-2-aminoethanesulfonic acid (TES), 10 trehalose, 10 glucose, 2 sucrose, 26 NaHCO_3_, 1 NaH_2_PO_4_, 1.5 CaCl_2_, 4 MgCl_2_. All data were digitized by a Digidata 1550b (Molecular Devices) at 10 kHz, except for the two-photon images, which were acquired using PrairieView (Bruker) at varying frequencies and saved as tiff files for later analysis. We used the frame triggers associated with our imaging frames (from Prairie View), recorded on Digidata 1550b, to carefully align odorant delivery with Ca^2+^ imaging measurements.

#### Image registration

The image stacks were motion-corrected using non-rigid motion correction (NoRMCorre^39^) and then manually validated to check for motion artifacts.

#### Defining regions of interest

To analyze Ca^2+^ imaging data, we defined regions of interest (ROIs) in Fiji and Python for GCaMP signals from PN dendrites in one hemisphere, or both hemispheres when the PN dendritic signals are available. We treated the entire PN dendrites from one hemisphere as one ROI.

#### Calculating fluorescence intensities

We used ROIs, defined above, as the unit for calculating fluorescent intensities (see above). For each ROI, we calculated the mean pixel value at each time point and then used the method ΔF/F0 to calculate, where F0 is the mean of the lowest 5% of raw fluorescence values in a given ROI over time and ΔF is F – F0.

### Courtship assay

Flies were collected shortly after eclosion. Male flies were housed individually, whereas female flies (*Canton-S*) were housed in groups of approximately 10. All females used as courtship targets were 3–5 days old virgins. All males tested in the experiments had not mated. Males were 4–7 days old in Fig. 4 and Extended Data Fig. 8a–d and were 2 days old in Extended Data Fig. 8e–i to lower the courtship baseline in males. All male flies were either *w^+^*, or *w^−^* but carried more than three mini-white markers from the transgenes they possessed. In single-pair courtship assays, two males (or one male and one female) were introduced into a custom-made courtship chamber with a diameter of 2 cm. In the courtship chain assay, five DA1-ORNs→VA1v-PNs males were introduced into a custom-made courtship chamber with a diameter of 5 cm. Courtship experiments were conducted under low white light to reduce baseline courtship activity, as vision is well known to influence the vigor of fly courtship. Before being placed into the courtship chamber, flies were briefly grouped in a tube and anesthetized on ice for less than 10 seconds. Once placed into the chamber, most flies were able to move immediately but did not fly away. Fly behavior was recorded for >25 minutes with a video camera at 13 frames per second and the first 25 minutes were quantified. In the single-pair male-male courtship assay, a control male and a rewired male were age-matched, and one of them was marked with an oil paint marker (Sharpie) on their thorax at least one day before the experiment. The paint was alternated between control and rewired males. 660nm LED lights were used to activate DA1-ORNs expressing csChrimson.

### Data and materials availability

All data are included in the manuscript, the supplementary materials, or are available upon request.

## Code availability

Code are available on github (https://github.com/Cheng-Lyu/CL_Stanford.git).

## Acknowledgement

We thank the Bloomington *Drosophila* Stock Center, the Vienna *Drosophila* Resource Center, and the *Drosophila* Genetic Resource Consortium for fly stocks, Gaby Maimon for custom plates and Tom Clandinin for extracellular saline used in Ca^2+^ imaging, Heather Dionne and Gerry Rubin for enhancer plasmids, Lorna Jayne for assistance with some VA1d-ORN experiments, and Tom Clandinin, Kang Shen, Gaby Maimon, Wei Qi, Sachin Sethi, as well as members of the Luo laboratory, especially Tom Hindmarsh Sten and Dan Pederick, for helpful discussions. This work was supported by National Institutes of Health grant (R01-DC005982 to LL). CL was supported by the Stanford Science Fellows Program. LL is an investigator of Howard Hughes Medical Institute.

## Author contributions

CL and LL conceived of the project. CL performed all of the experiments and analyzed the data. CL, ZL and LL jointly interpreted the data and decided on new experiments. ZL and CX assisted in cloning. JK assisted in behavioral experiments. CL and LL wrote the paper, with inputs from all other co-authors. LL supervised the work.

---

**Extended Data Fig. 1.**
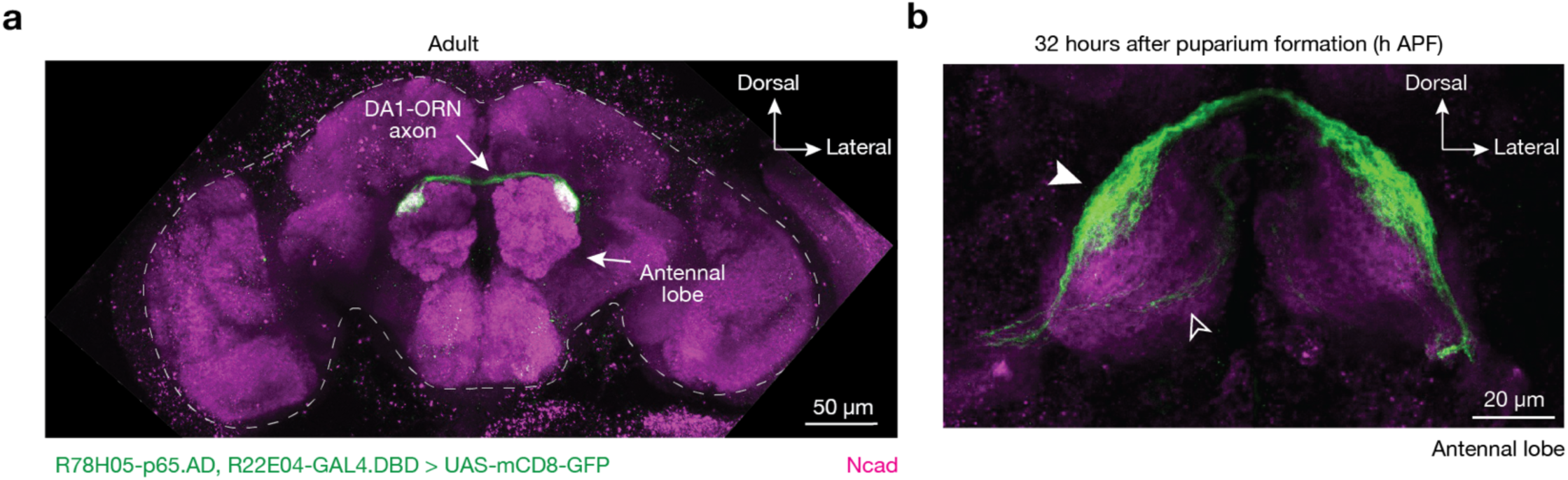
DA1-ORN split-GAL4 characterization. **a**, In adults, the split-GAL4, R78H05-AD + R22E04-DBD, labels DA1-ORNs in the whole brain as revealed by GFP staining (green). The brain border is dash-outlined. **b**, At around 32h APF, the same split-GAL4 driver most strongly labels DA1-ORNs (solid arrowhead) and weakly and sparsely labels a few other ORN types whose axons take the ventromedial trajectory (open arrowhead). Magenta: N-cadherin (Ncad) staining for neuropils. Maximum z-projections are shown.

**Extended Data Fig. 2.**
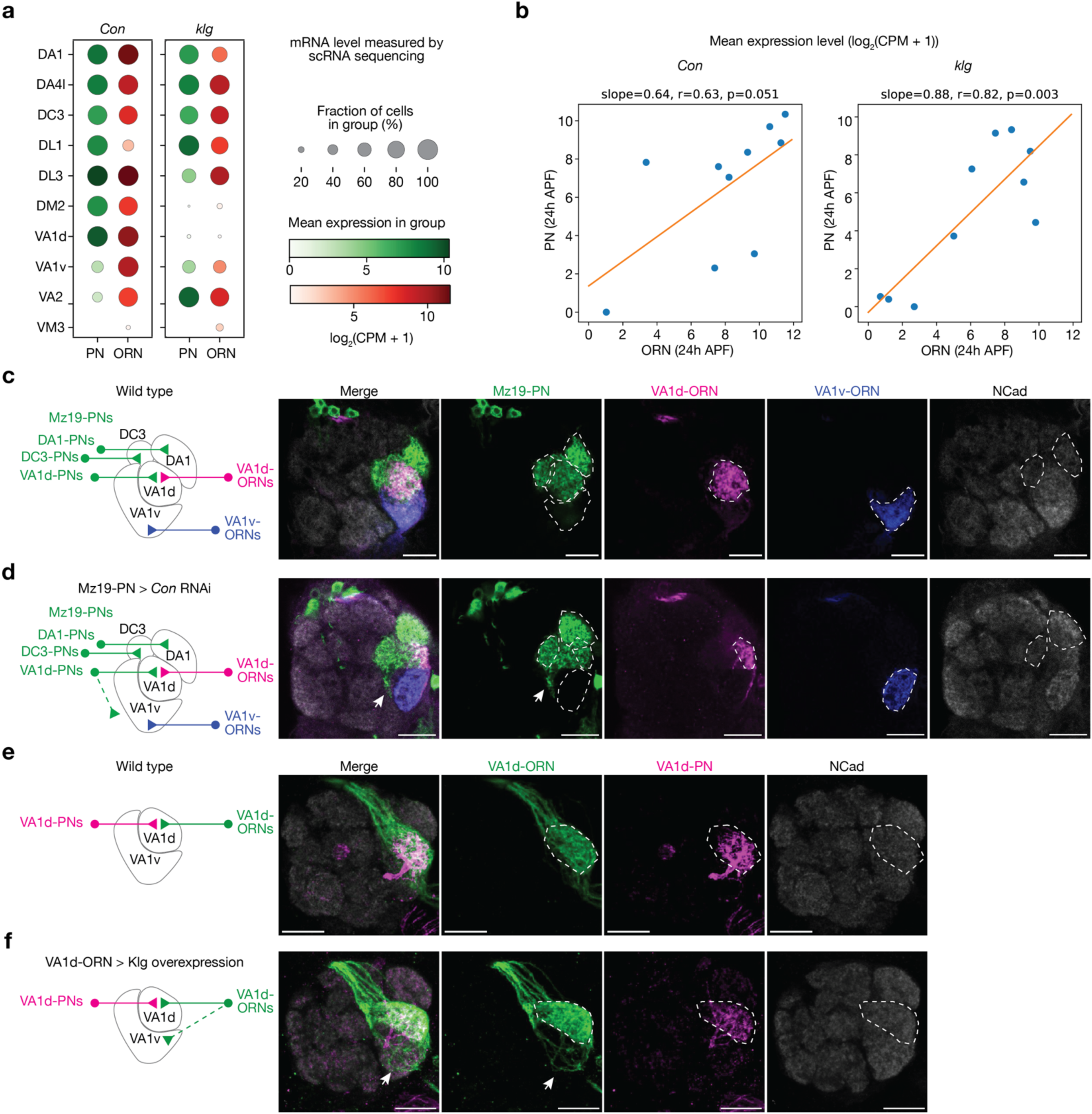
Cell-surface proteins Klg and Con regulate synaptic partner matching in the fly olfactory circuit. Connectin (Con) and Klingon (Klg) are homophilic adhesion molecules and have previously been reported to regulate the wiring of the *Drosophila* visual circuit and neuromuscular system, respectively^17,18^. Here, both the expression pattern and the genetic manipulation results suggest Con and Klg regulate synaptic partner matching of the *Drosophila* olfactory circuits. **a, b,** At around 24 hours after puparium formation (h APF), Con and Klg exhibit matching expression patterns across ORN and PN types based on single-cell RNA-sequencing (scRNAseq) data^19,20^. Dot plot (**a**) and scatter plot (**b**) with linear fitting (orange solid line) are shown. Blue dots in **b** represent the glomerular types shown in (**a**). **c**, Maximum projection of optical sections of the same antennal lobes from a wild-type brain. DA1-PNs, DC3-PNs, and VA1d-PNs (green) are labeled by GFP using *Mz19-GAL4*. VA1d-ORNs (magenta) are labeled by tdTomato using the *Or88a* promoter. VA1v-ORNs (blue) are labeled by rat CD2 using the *Or47b* promoter. The borders of DA1 and DC3 glomeruli are outlined based on the N-cadherin (NCad) staining signal. The VA1v and VA1d glomeruli are outlined according to the ORN signals. The glomerular targeting of these neurons in the wild-type brain is summarized by the schematic on the left. **d**, Same as **c**, but expressing *con* RNAi in the three Mz19+ PN types. The dendrites of some PNs, likely VA1d-PNs based on anatomical tracing, ectopically target outside glomeruli (arrows). 14 out 14 antennal lobes show similar phenotype using RNAi line 17898 from Vienna *Drosophila* Resource Center and 6 out of 14 antennal lobes show similar phenotype using RNAi line 28967 from Bloomington *Drosophila* Stock Center. **e**, Maximum projection of optical sections of the same antennal lobe from a wild-type brain, with VA1d-ORNs labeled using GFP (green) by GAL4/UAS and VA1d-PNs labeled using tdTomato (magenta) by QF2/QUAS. The border of VA1d glomerulus is outlined based on the N-cadherin (NCad) staining signal. The VA1d-ORN axons and VA1d-PN dendrites match well. **f**, Same as **e**, but overexpressing Klg in VA1d-ORNs. The axons of some VA1d-ORNs ectopically mistarget to the VA1v glomerulus (arrows).

**Extended Data Fig. 3.**
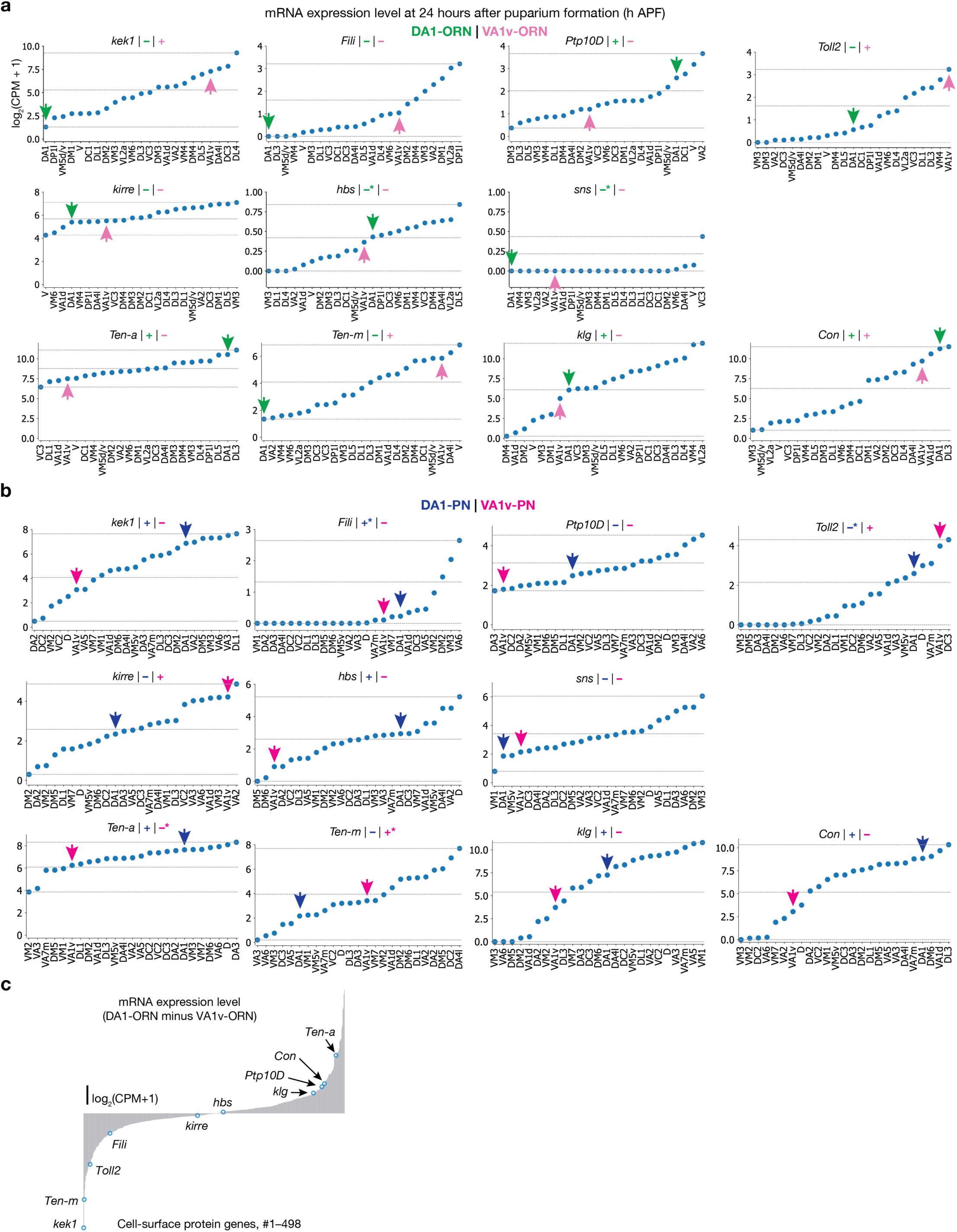
Expression levels of the CSPs in the developing *Drosophila* olfactory system, used in the DA1-ORN rewiring experiments. Here we provide the basis of assigning “+” or “–” for the expression levels of CSPs in Fig. 1d. **a**, mRNA expression levels of the wiring molecules used in the DA1-ORN rewiring experiments. The expression levels are based on the single-cell RNA sequencing (scRNAseq) data^19,20^ and all the ORN types decoded are shown. Plots are generated using data at ∼24 hours after puparium formation (h APF) in this and all other panels. In each subplot, dashed horizontal lines represent the lowest, the highest, and the median (the average of the minimum and maximum) expression levels. The green arrow indicates the data from DA1-ORNs, and the magenta arrow indicates the data from VA1v-ORNs. The ‘+’ or ‘–’ sign indicates the expression level as inferred from the scRNAseq data based on whether the expression level is above (‘+’) or below (‘–’) the median. Since the scRNAseq data are prone to measurement noise and may not accurately reflect protein expression due to post-transcriptional regulation, we corrected expression levels using the protein data and *in vivo* genetic manipulation results in CSPs where additional data were available. * designates places where corrections about the ‘+’ or ‘–’ are made, and the sign showed here is after the correction. Hbs and Sns are considered lowly expressed in both ORN types because of the absolute expression levels of these two mRNAs are very low. The unit of the y axis is log_2_(counts per million read + 1) in this and all other panels. **b**, Same as **a**, but plotting the expression level in all the PN types decoded. The blue arrow indicates the data from DA1-PNs and the magenta arrow indicates the data from VA1v-PNs. Fili is considered highly expressed in DA1-PNs based on the data from a previous study (in Fig. 3D)^9^. Toll2 is considered lowly expressed in DA1-PNs based on the conditional-tag data from the companion study (Fig. 1d)^5^. Ten-a is considered lowly expressed in VA1v-PNs and Ten-m is considered highly expressed in VA1v-PNs based on the antibody staining data from a previous study (Fig. 2)^6^. **c**, The difference of mRNA expression level of CSPs that are expressed between DA1-ORNs and VA1v-ORNs, with the 10 wiring molecules used in the rewiring experiments indicated. *Sticks and stones* (*sns*), encoding a second interacting partner of kirre in addition to Hbs (companion study^5^), is not included here since it is lowly expressed in both ORN types.

**Extended Data Fig. 4.**
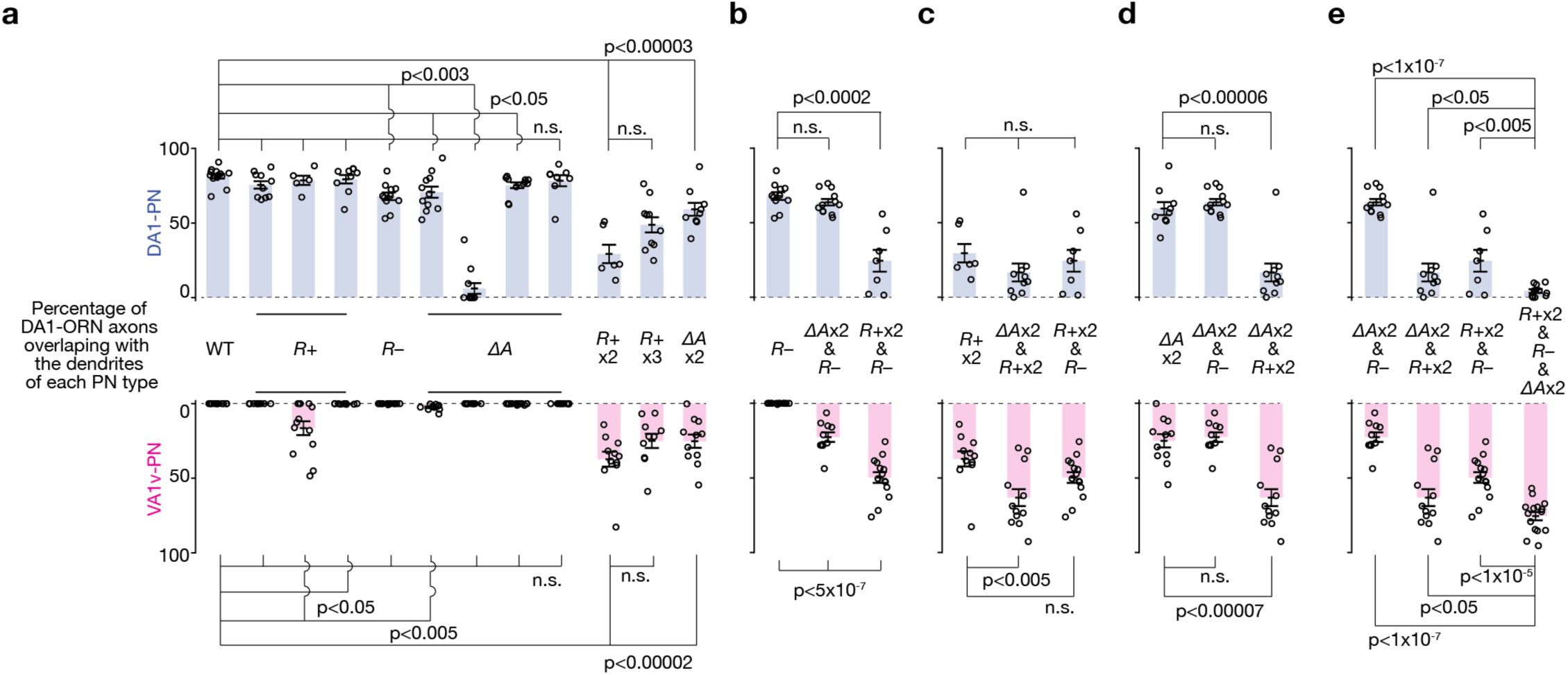
Statistical tests in the DA1-ORNs®VA1v-PNs rewiring. Same plots as in Fig. 2a, but with statistical tests added. Percentage of DA1-ORN axons overlapping with the dendrites of DA1-PNs (top) and VA1v-PNs (bottom). Circles indicate individual antennal lobes; bars indicate the population mean ± s.e.m. ‘*R+*’: Kek1 overexpression (OE), Fili OE, and Kirre OE (left to right). ‘*R–*’: *Ptp10D* RNAi. ‘*ΔA*’: *ten-a* RNAi, Ten-m OE, *klg* RNAi, and *con* RNAi (left to right). ‘*R+* x2’: Kek1 OE + Fili OE. ‘*R+* x3’: Kek1 OE + Fili OE + Kirre OE. ‘*ΔA* x2’: *ten-a* RNAi + *con* RNAi. Unpaired t-test was used. **a**, All statistical tests were performed between ‘WT’ and an individual manipulation condition, respectively, except the one test where it is performed between ‘*R+* x2’ and ‘*R+* x3’. In all the eight single-CSP manipulations, six—Fili OE, Kirre OE, *Ptp10D* RNAi, *ten-a* RNAi, Ten-m OE, and *klg* RNAi—showed observable yet very mild DA1-ORNs®VA1v-PNs mismatching phenotypes, with some reaching statistical significance while others not (n.s.). **b**, All tests were performed between ‘*R–*’ and other manipulation conditions, respectively. **c**, All tests were performed between ‘*R+* x2’ and other manipulation conditions, respectively. **d**, All tests were performed between ‘*ΔA* x2’ and other manipulation conditions, respectively. **e**, All tests were performed between ‘*R+* x2 & *R–* & *ΔA* x2’ (the DA1-ORN rewired condition) and other manipulation conditions, respectively.

**Extended Data Fig. 5.**
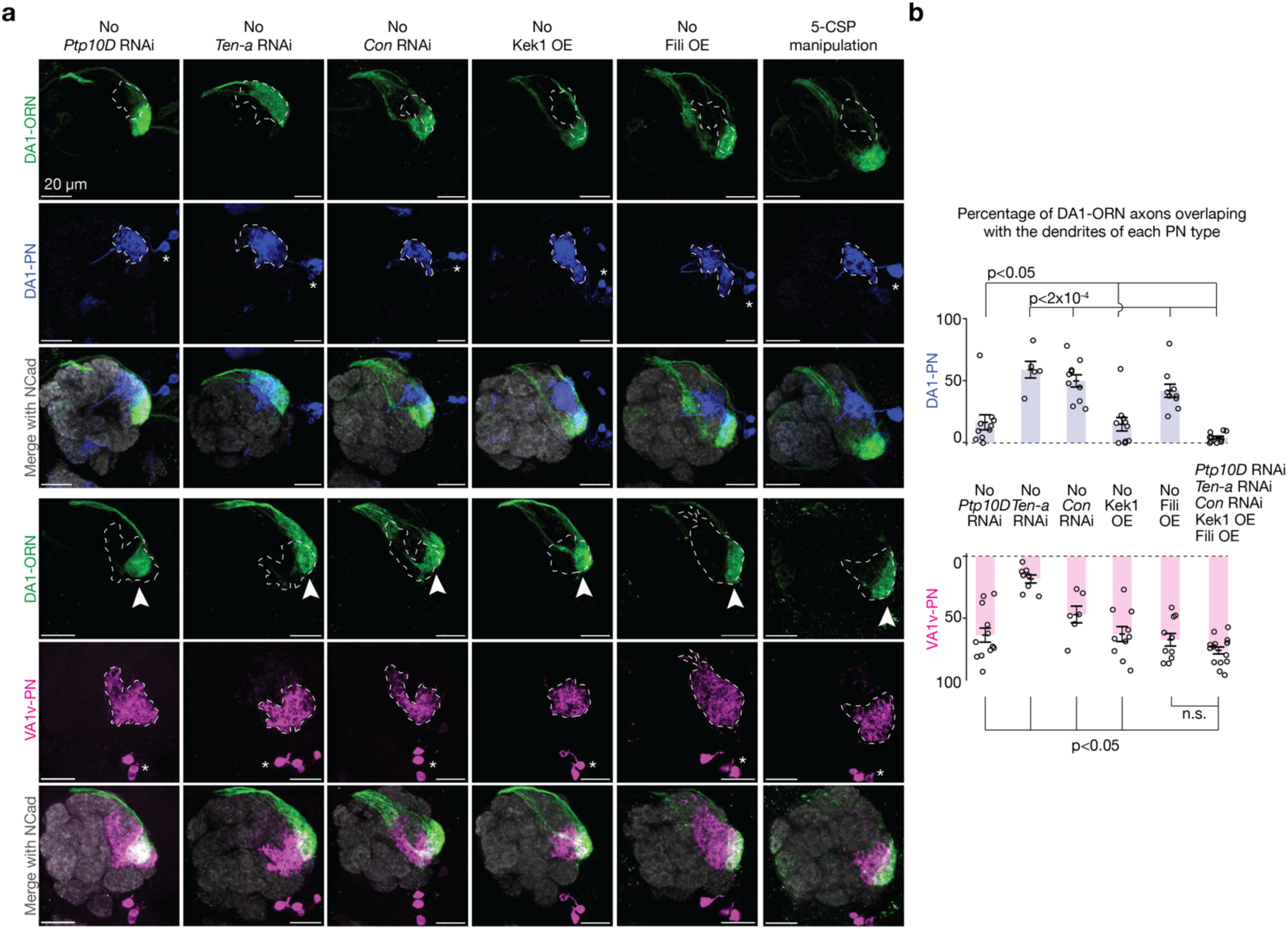
Omitting any one of the five CSP changes reduced the magnitude of DA1-ORN rewiring. **a**, Genetic manipulations are labeled on the top. Maximum z-projections of adult antennal lobes around DA1-ORN axons (green) are shown. Top three rows: DA1-PNs (blue) are co-labeled with borders dashed outlined. Bottom three rows: VA1v-PNs (magenta) are co-labeled with borders dashed outlined. The genetic manipulation condition for the rightmost column contains five genetic changes as listed, same as the rewiring condition as in Fig. 2c. Genetic manipulation conditions for the left five columns are the same as for the rightmost column except missing one manipulation as indicated. Arrowheads indicate the mismatch of DA1-ORN axons with VA1v-PN dendrites; scale bar = 20 µm; * designates PN cell bodies. **b**, Percentage of DA1-ORN axons overlapping with the dendrites of DA1-PNs (top) and VA1v-PNs (bottom). Same genetic manipulation conditions as **a**. Circles indicate individual antennal lobes; bars indicate the population mean ± s.e.m. All the tests are performed between the data from the rightmost column and the data from other columns, respectively. Unpaired t-test is used. n.s., not significant.

**Extended Data Fig. 6.**
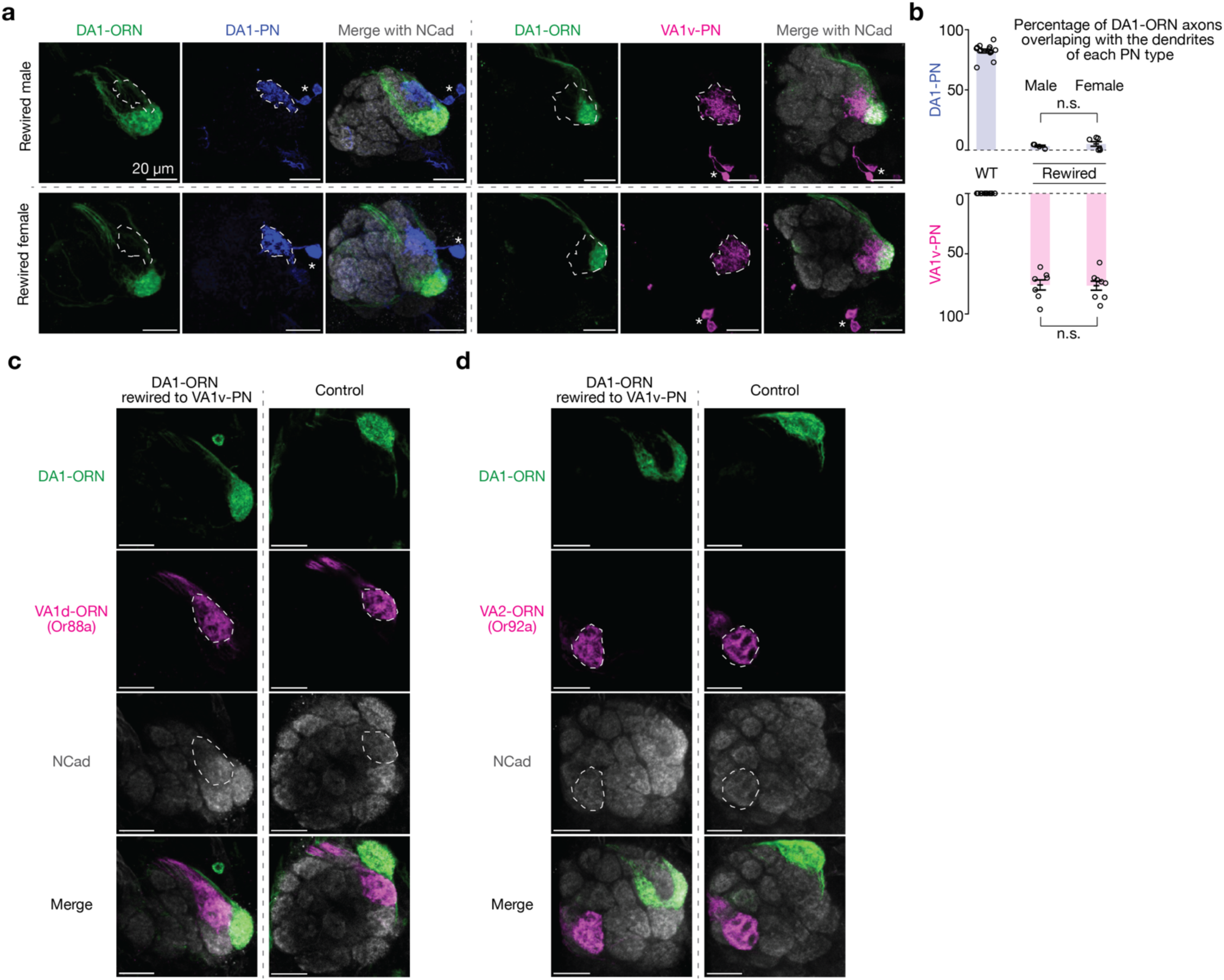
Males and females exhibit similar DA1-ORN→VA1v-PN rewiring and the axons of some non-DA1/non-VA1v ORNs remain confined in their original glomeruli. **a**, Maximum z-projections of adult antennal lobes around DA1-ORN axons (green) are shown. Top row: rewired male; bottom row: rewired female. Left three columns: DA1-PNs (blue) are co-labeled with borders dashed outlined; right three columns: VA1v-PNs (magenta) are co-labeled with borders dashed outlined. The genetic manipulation condition is the same as the rewiring condition as in Fig. 2c. Scale bar = 20 µm here and throughout the figure; * designates PN cell bodies. **b**, Percentage of DA1-ORN axons overlapping with the dendrites of DA1-PNs (top) and VA1v-PNs (bottom). Circles indicate individual antennal lobes; bars indicate the population mean ± s.e.m. Unpaired t-tests are performed. **c**, Here we tested whether in DA1-ORN→VA1v-PN rewired flies, VA1d-ORN axons target normally to the VA1d glomerulus, which is in between DA1 and VA1v. Shown are maximum z-projection of adult antennal lobes around DA1-ORN (green, labeled by a membrane-targeted GFP driven by a split-GAL4) and VA1d-ORN (magenta, *Or88a* promotor driving tdTomato) axons. Gray: N-cadherin (Ncad) staining for neuropils. Scale bar = 20 µm. The VA1d glomerulus boarders are dash-outlined based on VA1d-ORN signals. The axons of VA1d-ORNs remain confined to their endogenous glomerulus. Note ventral shift of DA1-ORN axon terminals in rewired flies compared with control, consistent with their matching with VA1v-PN dendrites. **d**, Same as **c**, but with VA2-ORN co-labeled using Or92a promotor driving rat CD2, showing that they are confined within the VA2 glomerulus (outlined) in the rewired fly as in control.

**Extended Data Fig. 7.**
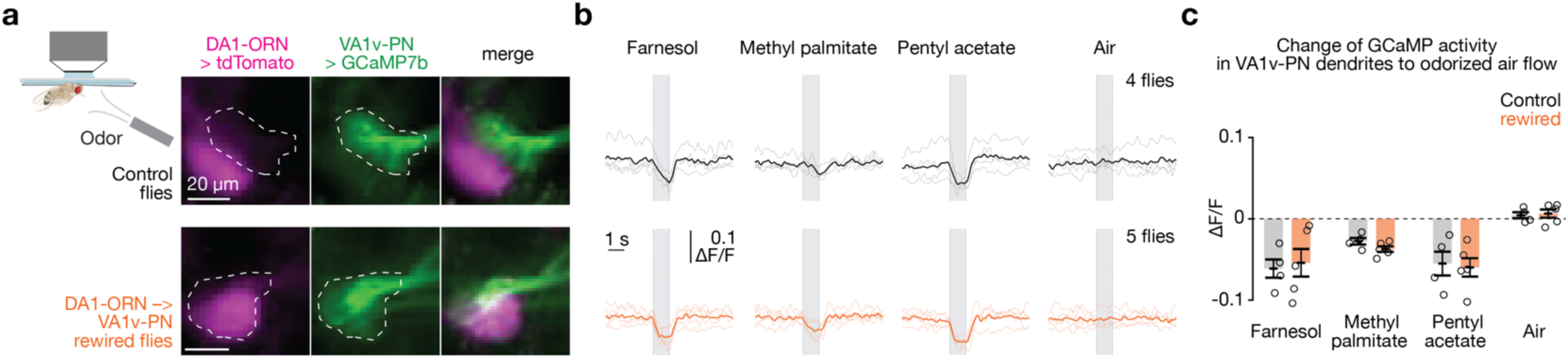
VA1v-PNs in DA1-ORN rewired flies retain similar levels of inhibitory response to non-cognate odors tested. **a**, Imaging neural activity in a plate-tethered fly with odorized air flow delivered to the fly antennae. Images of tdTomato signal in DA1-ORN axons and GCaMP7b signal in VA1v-PN dendrites are shown from a control fly (top) and a DA1-ORN rewired fly (bottom). Images are averaged across the entire recording. The VA1v glomerulus is outlined according to GCaMP7b signal. Scale bar = 20 µm. **b**, Averaged GCaMP7b activity in VA1v-PN dendrites in response to odorized air flows, measured by fluorescence intensity change over baseline (ΔF/F). Top: control flies. Bottom: DA1-ORN rewired flies. The grey vertical stripes indicate odorized air flows (1 s each). Light-colored traces indicate the means of individual flies; dark-colored traces indicate the population mean. Farnesol strongly activates DC3-ORNs^36^ (spatially close to DA1 and VA1v glomeruli), fly pheromone methyl palmitate mainly activates VA1d-ORNs^28^ (in between DA1 and VA1v glomeruli), and pentyl acetate activates a variety types of ORNs^40^. **c**, Change of GCaMP7b activity in VA1v-PN dendrites to odorized air flows. The change of activity is calculated by subtracting the average GCaMP7b activity in the 0.5 s before the onset of odor delivery from that in the 0.4 s flanking the offset of odor delivery. Circles indicate the means of individual flies; bars indicate the population mean ± s.e.m. VA1v-PNs in DA1-ORN rewired flies retain similar levels of inhibitory response to non-cognate odors tested as in wild-type flies. Note that the inhibitory response to MP is weaker than the response to other two odors, consistent with the low volatility of the large molecular weight of MP. P < 0.02 for all odorized responses to the control air group. Unpaired t-tests were used.

**Extended Data Fig. 8.**
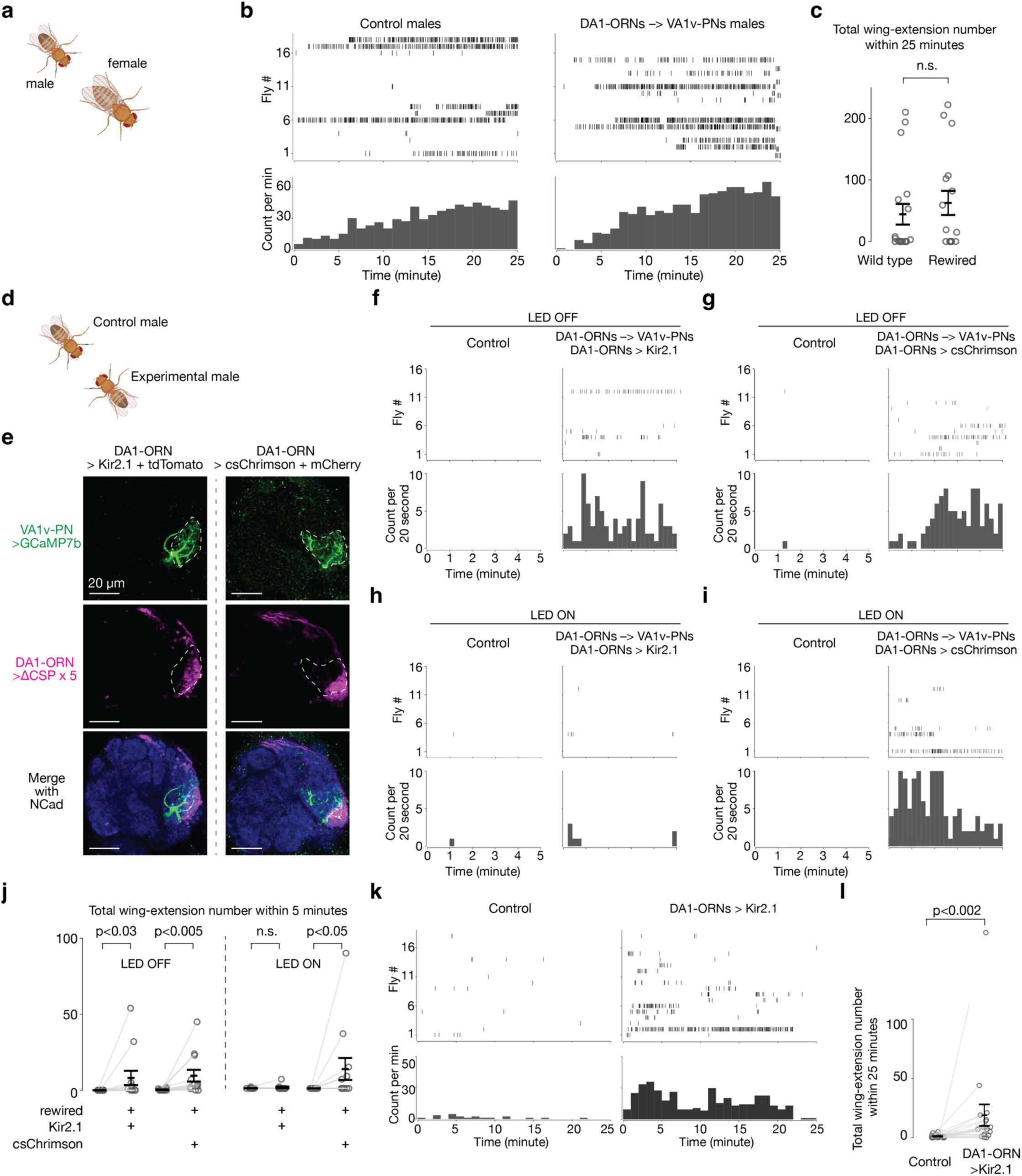
Additional behavioral examination of the courtship activity of DA1-ORN→VA1v-PN rewired males. **a–c**, Wild-type and DA1-ORN→VA1v-PN rewired males exhibit similar courtship activity towards virgin females. **a**, Courtship assay where one male and one virgin female are introduced in the same behavioral chamber to monitor the courtship activity from the male towards the female. The chamber diameter is 2 cm. **b**, Unilateral-wing-extension rasters (top) and extension count per minute (1-min bins, bottom). Left: wild-type males; right: DA1-ORN rewired males. **c**, Total wing-extension number during the 25-minute recordings. Circles indicate individual flies; bars indicate the population mean ± s.e.m. Wilcoxon signed-rank test is used given the non-normal distribution of the data points. **d–l,** In DA1-ORN→VA1v-PN rewired males, two connectivity changes could potentially contribute to behavioral changes: (1) DA1-ORNs losing endogenous connection to DA1-PNs (loss-of-connection or LoC) and (2) DA1-ORNs gain new connection to VA1v-PNs (gain-of-connection or GoC). Two experiments here tested if both LoC and GoC contribute to the increased male-male courtship in rewired flies. **d**, Courtship assay where one wild-type male and one experimental male are introduced in the same behavioral chamber to monitor their courtship activity towards each other. Two-day old males are used here and throughout the figure to lower the courtship baseline in males. **e**, Maximum z-projection of adult antennal lobes around DA1-ORN axons (magenta, labeled by a membrane-targeted tdTomato driven by a split-GAL4). DA1-PN dendrites are co-labeled (green, labeled by GCaMP driven by a LexA driver) with borders dash-outlined. Scale bar = 20 µm. DA1-ORNs also express the same 5 cell-surface-proteins (CSP) as the rewired flies in Fig. 2d. The rewiring of DA1-ORN axons to VA1v-PN dendrites still persist with exogenously expressing csChrimson (a red-shifted channelrhodopsin for activating neurons when LED is on) or Kir2.1 (an inward rectifying K^+^ channel for silencing neuronal activity) across development. **f**, Same as **b**, but for the male-male courtship assay where one male is the control and the other male has DA1-ORNs rewired to VA1v-PNs and silenced by additional expression of Kir2.1. The recording duration was reduced to 5 minutes. Red LEDs are turned off. **g**, Same as **f**, but the experimental male is changed to one with DA1-ORNs rewired to VA1v-PNs and exogenously expressing csChrimson. **h, i**, Same as **f, g**, but with LEDs turned on to activate the males expressing csChrimson. **j**, Total wing-extension number during the 5-minute recordings. Circles indicate individual flies; bars indicate the population mean ± s.e.m. Wilcoxon signed-rank test is used given the non-normal distribution of the data points. **k, l**, Same as **b, c**, but for the male-male courtship assay where one male is the control and the other male with DA1-ORNs silenced by exogenously expressing Kir2.1. (1) To test the contribution LoC, we compared the courtship activity of wild-type males versus DA1-ORN rewired but silenced (by exogenously expressing inward-rectifier potassium channel Kir2.1) males. If LoC contributed to increased courtship activity in rewired males, we should expect to see DA1-ORN rewired but silenced males show stronger courtship activity than wild-type males. Results (**f, j**) support this working model. This result is not surprising, since silencing DA1-ORNs alone yields similar behavioral phenotype (**k**, **l** and ref^13^ using Or67d mutant). (2) To test the contribution of GoC, we note that DA1-ORN rewired flies simultaneously have LoC. Therefore, we sought to compare the difference between flies with ‘GoC & LoC’ (males with DA1-ORN rewired and can be optogenetically activated) and flies with ‘LoC’ alone (males with DA1-ORN rewired but silenced). Stronger courtship activity in flies with ‘GoC & LoC’ would supports that ‘GoC’ has a positive contribution on rewired male’s enhanced courtship activity. Results (**f–j**) support this working model. When LEDs were turned off, rewired males from both groups showed stronger courtship activity over wild-type males, supporting that ‘LoC’ alone could increase male courtship activity towards other males. When red LEDs were turned on, the courtship difference in ‘LoC’ group disappeared. Since neither male from this group exogenously expressed any light-sensitive channels, we hypothesized that bright lighting might inhibit male courtship activity. However, in males with DA1-ORN rewired and exogenously expressing csChrimson, they exhibited significantly stronger courtship activity over wild-type males despite the lighting effect. This supports that activation of DA1-ORNs in rewired males contribute to the increase of male-male courtship activity.

**Extended Data Fig. 9.**
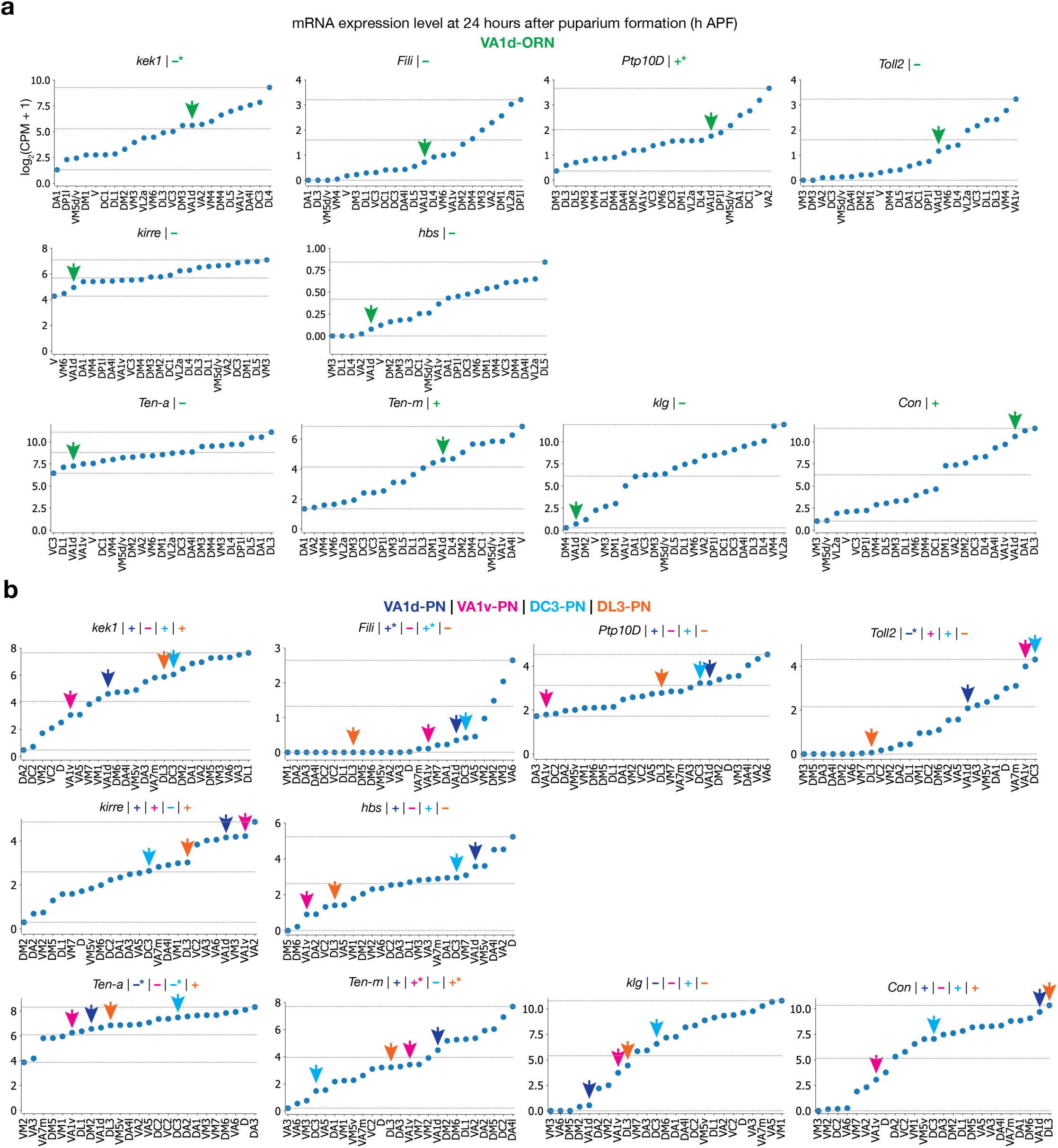
Expression levels of the ten CSPs in the VA1d-ORN rewiring. Here we provide the basis of assigning “+” or “–” for the expression levels of CSPs in Fig. 5a. **a**, mRNA expression levels of the 10 CSPs used in the VA1d-ORN rewiring experiments. The expression levels are based on the single-cell RNA-sequencing (scRNAseq) data^19,20^ and all the ORN types decoded are shown. Plots are generated using data at 24 hours after puparium formation (h APF) in this and all other panels. In each subplot, the lowest and highest expression levels are indicated by the dashed horizontal lines. The green arrow indicates the data from VA1d-ORNs. The ‘+’ or ‘–’ sign indicates the expression level as inferred from the scRNAseq data based on whether the expression level is above (‘+’) or below (‘–’) the median, which is the average of the minimum and maximum value. Since the scRNAseq data are prone to measurement noise and may not accurately reflect protein expression due to post-transcriptional regulation, we corrected the RNA data using the protein data and the *in vivo* genetic manipulation results in CSPs where additional data were available. * designates places where corrections are made, and the sign shown here is after the correction. Kek1 is considered lowly expressed based on the conditional-tag data from the companion study (Extended Data Fig. 2)^5^. Fili is considered highly expressed in DA1-PNs based on the data from a previous study (in Fig. 3D)^9^. Because the RNA level in DC3-PNs is higher than in DA1-PNs, we considered Fili also highly expressed in DC3-PNs. Ptp10D is considered highly expressed based on the conditional-tag data as well as the knockout experiments from the companion study (Figs. 1c and 2c)^5^. The unit of the y axis is log_2_(counts per million read + 1). **b**, Same as **a**, but plotting the expression level in all the PN types decoded. The blue arrow indicates the data from DA1-PNs, the magenta arrow indicates the data from VA1v-PNs, light blue arrow indicates the data from DC3-PNs, and the orange arrow indicates the data from DL3-PNs. Toll2 is considered lowly expressed in VA1d-PNs based on the conditional-tag data from the companion study (Fig. 1d)^5^. Ten-a is considered lowly expressed in VA1d- and DL3-PNs, whereas Ten-m is considered highly expressed in VA1v- and DL3-PNs based on the antibody staining data from a previous study (Fig. 2)^6^.

**Extended Data Fig. 10.**
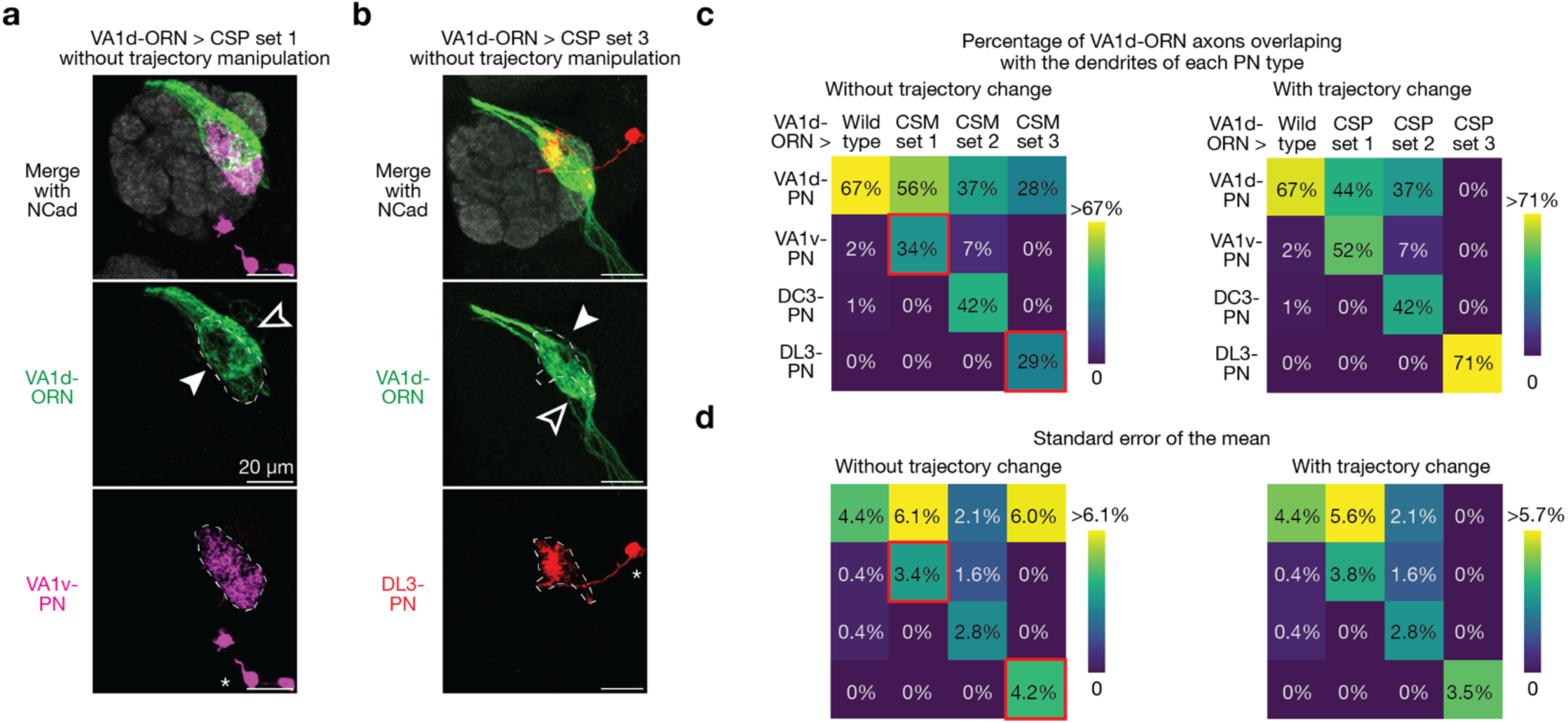
Trajectory manipulation of VA1d-ORN axons is necessary for rewiring with the dendrites of VA1v- and DL3-PNs. **a**, Maximum z-projections of adult antennal lobes around VA1d-ORN axons in the rewiring experiments without trajectory manipulation of VA1d-ORN axons. Genetic manipulations = Kek1 overexpression (OE) + *Ptp10D* RNAi + *con* RNAi. VA1v-PNs (magenta) are co-labeled with borders dash-outlined. The solid arrowhead indicates the mismatch of VA1d-ORN axons with VA1v-PN dendrites; the open arrowhead indicates the part of VA1d-ORN axons that do not match with VA1v-PN dendrites. Scale bar = 20 µm; * designates PN cell bodies. **b**, Same as **a**, but for the rewiring experiment of VA1d-ORN axons to DL3-PN dendrites. Genetic manipulations = Ten-a OE. DL3-PNs (red) are co-labeled with borders dash-outlined. The solid arrowhead indicates the mismatch of VA1d-ORN axons with DL3-PN dendrites; the open arrowhead indicates the part of VA1d-ORN axons that do not match with DL3-PN dendrites. **c**, Percentage of VA1d-ORN axons overlapping with the dendrites of each PN type (indicated on the left) in wild type and the three rewired conditions without (left) and with (right) trajectory manipulations. The right matrix is a repeat of Fig. 5c for ease of comparison. The two red boxes in the left matrix indicate the genetic manipulation conditions showed in (**a**) and (**b**), respectively. Note that in these two boxes, the ratios of VA1d-ORN axons rewired to the target PNs are less compared to the conditions with trajectory manipulations, suggesting that trajectory manipulation of VA1d-ORN axons is necessary in the rewiring to the dendrites of VA1v- and DL3-PNs. n ≥ 6 for all the conditions. **d**, Same as **c**, but plotting the s.e.m. instead of the population mean.

**Extended Data Table 1.** Summary of genotypes used in each experiment, arranged according to figure panels.

| Figure and panel | Brief description | Fly genotype |  |  |
| --- | --- | --- | --- | --- |
|  |  | x chromosome | 2nd chromosome | 3rd chromosome |
| **Figure 1** |  |  |  |  |
| b | wild type DA1-ORN and DA1-PN | *UAS-Dcr2* (BDSC 24646), *UAS-mCD8-GFP* (BDSC 24648) / *w or y* | *R24A10-QF2* (this work), *QUAS-mtdTomato-HA* (BDSC 30004) / + | *R78H05-p65.AD* (BDSC 601815), *R22E04-GAL4.DBD* (BDSC 68303) / + |
| c | wild type DA1-ORN and VA1v-PN | *UAS-Dcr2* / *w or y* | *VT003280-LexA* (BDSC 94678), *lexAop-rCD2::RFP-p10.UAS-mCD8::GFP-p10* (BDSC 67093) / + | *R78H05-p65.AD*, *R22E04-GAL4.DBD* / + |
| h | Kek1 overexpression with DA1-PN colabeling | *UAS-Dcr2*, *UAS-mCD8-GFP* / *w or y* | *R24A10-QF2*, *QUAS-mtdTomato-HA* / *UAS-Kek1* (BDSC 43665) | *R78H05-p65.AD*, *R22E04-GAL4.DBD* / + |
|  | Fili overexpression with DA1-PN colabeling | *UAS-Dcr2*, *UAS-mCD8-GFP* / *w or y* | *R24A10-QF2*, *QUAS-mtdTomato-HA* / + | *R78H05-p65.AD*, *R22E04-GAL4.DBD* / *UAS-Fili* (DGRC 109636) |
|  | Kirre overexpression with DA1-PN colabeling | *UAS-Dcr2*, *UAS-mCD8-GFP* / *UAS-Kirre* (BDSC 26609) | *R24A10-QF2*, *QUAS-mtdTomato-HA* / + | *R78H05-p65.AD*, *R22E04-GAL4.DBD* / + |
|  | Ptp10D RNAi with DA1-PN colabeling | *UAS-Dcr2*, *UAS-mCD8-GFP* / *w or y* | *R24A10-QF2*, *QUAS-mtdTomato-HA* / + | *R78H05-p65.AD*, *R22E04-GAL4.DBD* / *UAS-Ptp10D-RNAi* (VDRC 8010) |
|  | *ten-a* RNAi with DA1-PN colabeling | *UAS-Dcr2*, *UAS-mCD8-GFP* / *w or y* | *R24A10-QF2*, *QUAS-mtdTomato-HA* / + | *R78H05-p65.AD*, *R22E04-GAL4.DBD* / *UAS-Ten-a-RNAi* (BDSC 29439) |
|  | Ten-m overexpression with DA1-PN colabeling | *UAS-Dcr2*, *UAS-mCD8-GFP* / *w or y* | *R24A10-QF2*, *QUAS-mtdTomato-HA* / + | *R78H05-p65.AD*, *R22E04-GAL4.DBD* / *UAS-Ten-m* (BDSC 41567) |
|  | *klg* RNAi with DA1-PN colabeling | *UAS-Dcr2*, *UAS-mCD8-GFP* / *w or y* | *R24A10-QF2*, *QUAS-mtdTomato-HA* / + | *R78H05-p65.AD*, *R22E04-GAL4.DBD* / *UAS-klg-RNAi* (BDSC 28746) |
|  | *con* RNAi with DA1-PN colabeling | *UAS-Dcr2*, *UAS-mCD8-GFP* / *w or y* | *R24A10-QF2*, *QUAS-mtdTomato-HA* / *UAS-con-RNAi* (VDRC 17898) | *R78H05-p65.AD*, *R22E04-GAL4.DBD* / + |
|  | Kek1 overexpression with VA1v-PN colabeling | *UAS-Dcr2* / *w or y* | *VT003280-LexA*, *lexAop-rCD2::RFP-p10.UAS-mCD8::GFP-p10* / *UAS-Kek1* | *R78H05-p65.AD*, *R22E04-GAL4.DBD* / + |
|  | Fili overexpression with VA1v-PN colabeling | *UAS-Dcr2* / *w or y* | *VT003280-LexA*, *lexAop-rCD2::RFP-p10.UAS-mCD8::GFP-p10* / + | *R78H05-p65.AD*, *R22E04-GAL4.DBD* / *UAS-Fili* |
|  | Kirre overexpression with VA1v-PN colabeling | *UAS-Dcr2* / *UAS-Kirre* | *VT003280-LexA*, *lexAop-rCD2::RFP-p10.UAS-mCD8::GFP-p10* / + | *R78H05-p65.AD*, *R22E04-GAL4.DBD* / + |
|  | *Ptp10D* RNAi with VA1v-PN colabeling | *UAS-Dcr2* / *w or y* | *VT003280-LexA*, *lexAop-rCD2::RFP-p10.UAS-mCD8::GFP-p10* / + | *R78H05-p65.AD*, *R22E04-GAL4.DBD* / *UAS-Ptp10D-RNAi* |
|  | *ten-a* RNAi with VA1v-PN colabeling | *UAS-Dcr2* / *w or y* | *VT003280-LexA*, *lexAop-rCD2::RFP-p10.UAS-mCD8::GFP-p10* / + | *R78H05-p65.AD*, *R22E04-GAL4.DBD* / *UAS-Ten-a-RNAi* |
|  | Ten-m overexpression with VA1v-PN colabeling | *UAS-Dcr2* / *w or y* | *VT003280-LexA*, *lexAop-rCD2::RFP-p10.UAS-mCD8::GFP-p10* / + | *R78H05-p65.AD*, *R22E04-GAL4.DBD* / *UAS-Ten-m* |
|  | *klg* RNAi with VA1v-PN colabeling | *UAS-Dcr2* / *w or y* | *VT003280-LexA*, *lexAop-rCD2::RFP-p10.UAS-mCD8::GFP-p10* / + | *R78H05-p65.AD*, *R22E04-GAL4.DBD* / *UAS-klg-RNAi* |
|  | *con* RNAi with VA1v-PN colabeling | *UAS-Dcr2* / *w or y* | *VT003280-LexA*, *lexAop-rCD2::RFP-p10.UAS-mCD8::GFP-p10* / *UAS-con-RNAi* | *R78H05-p65.AD*, *R22E04-GAL4.DBD* / + |
| **Figure 2** |  |  |  |  |
| b | *R+* x2 with DA1-PN colabeling | *UAS-Dcr2*, *UAS-mCD8-GFP* / *w or y* | *R24A10-QF2*, *QUAS-mtdTomato-HA* / *UAS-Kek1* | *R78H05-p65.AD*, *R22E04-GAL4.DBD* / *UAS-Fili* |
|  | *R+* x3 with DA1-PN colabeling | *UAS-Dcr2*, *UAS-mCD8-GFP* / *UAS-Kirre* | *R24A10-QF2*, *QUAS-mtdTomato-HA* / *UAS-Kek1* | *R78H05-p65.AD*, *R22E04-GAL4.DBD* / *UAS-Fili* |
| | $\Delta A$ x2 with DA1-PN colabeling | *UAS-Dcr2*, *UAS-mCD8-GFP* / *w or y* | *R24A10-QF2*, *QUAS-mtdTomato-HA* / *UAS-con-RNAi* | *R78H05-p65.AD*, *R22E04-GAL4.DBD* / *UAS-ten-a-RNAi* |
| | $\Delta A$ x2 & *R-* with DA1-PN colabeling | *UAS-Dcr2*, *UAS-mCD8-GFP* / *w or y* | *R24A10-QF2*, *QUAS-mtdTomato-HA* / *UAS-con-RNAi* | *R78H05-p65.AD*, *R22E04-GAL4.DBD* / *UAS-ten-a-RNAi*, *UAS-Ptp10D-RNAi* |
| | $\Delta A$ x2 & *R+* x2 with DA1-PN colabeling | *UAS-Dcr2*, *UAS-mCD8-GFP* / *w or y* | *R24A10-QF2*, *QUAS-mtdTomato-HA* / *UAS-con-RNAi*, *UAS-Kek1* | *R78H05-p65.AD*, *R22E04-GAL4.DBD* / *UAS-ten-a-RNAi*, *UAS-Fili* |
|  | R+ x2 & R- with DA1-PN colabeling | *UAS-Dcr2, UAS-mCD8-GFP / w or y* | *R24A10-QF2, QUAS-mtdTomato-HA / UAS-Kek1* | *R78H05-p65.AD, R22E04-GAL4.DBD / UAS-Fili, UAS-Ptp10D-RNAi* |
|  | R+ x2 with VA1v-PN colabeling | *UAS-Dcr2, UAS-mCD8-GFP / w or y* | *VT003280-LexA, lexAop-rCD2::RFP-p10.UAS-mCD8::GFP-p10 / UAS-Kek1* | *R78H05-p65.AD, R22E04-GAL4.DBD / UAS-Fili* |
|  | R+ x3 with VA1v-PN colabeling | *UAS-Dcr2, UAS-mCD8-GFP / UAS-Kirre* | *VT003280-LexA, lexAop-rCD2::RFP-p10.UAS-mCD8::GFP-p10 / UAS-Kek1* | *R78H05-p65.AD, R22E04-GAL4.DBD / UAS-Fili* |
| | $\Delta A$ x2 with VA1v-PN colabeling | *UAS-Dcr2, UAS-mCD8-GFP / w or y* | *VT003280-LexA, lexAop-rCD2::RFP-p10.UAS-mCD8::GFP-p10 / UAS-con-RNAi* | *R78H05-p65.AD, R22E04-GAL4.DBD / UAS-ten-a-RNAi* |
| | $\Delta A$ x2 & R- with VA1v-PN colabeling | *UAS-Dcr2, UAS-mCD8-GFP / w or y* | *VT003280-LexA, lexAop-rCD2::RFP-p10.UAS-mCD8::GFP-p10 / UAS-con-RNAi* | *R78H05-p65.AD, R22E04-GAL4.DBD / UAS-ten-a-RNAi, UAS-Ptp10D-RNAi* |
| | $\Delta A$ x2 & R+ x2 with VA1v-PN colabeling | *UAS-Dcr2, UAS-mCD8-GFP / w or y* | *VT003280-LexA, lexAop-rCD2::RFP-p10.UAS-mCD8::GFP-p10 / UAS-con-RNAi, UAS-Kek1* | *R78H05-p65.AD, R22E04-GAL4.DBD / UAS-ten-a-RNAi, UAS-Fili* |
|  | R+ x2 & R- with VA1v-PN colabeling | *UAS-Dcr2, UAS-mCD8-GFP / w or y* | *VT003280-LexA, lexAop-rCD2::RFP-p10.UAS-mCD8::GFP-p10 / UAS-Kek1* | *R78H05-p65.AD, R22E04-GAL4.DBD / UAS-Fili, UAS-Ptp10D-RNAi* |
| c | R+ x2 & R- & $\Delta A$ x2 with DA1-PN colabeling | *UAS-Dcr2, UAS-mCD8-GFP / w or y* | *R24A10-QF2, QUAS-mtdTomato-HA / UAS-con-RNAi, UAS-Kek1* | *R78H05-p65.AD, R22E04-GAL4.DBD / UAS-ten-a-RNAi, UAS-Fili, UAS-Ptp10D-RNAi* |
| | R+ x2 & R- & $\Delta A$ x2 with VA1v-PN and VA1v-ORN colabeling | *UAS-Dcr2, UAS-mCD8-GFP / w or y* | *VT003280-LexA, lexAop-rCD2::RFP-p10.UAS-mCD8::GFP-p10 / UAS-con-RNAi, UAS-Kek1, OR47b-ratCD2 (BDSC 9916)* | *R78H05-p65.AD, R22E04-GAL4.DBD / UAS-ten-a-RNAi, UAS-Fili, UAS-Ptp10D-RNAi* |
| **Figure 3** |  |  |  |  |
|  | Control | *UAS-Dcr2 / w* | *VT003280-LexA, UAS-IVS-myr-tdtomato (BDSC 32222) / VT003280-LexA* | *R78H05-p65.AD, R22E04-GAL4.DBD, LexAop-IVS-jGCaMP7b (BDSC 80915) / +* |
|  | DA1-ORN rewired | *UAS-Dcr2 / w* | *VT003280-LexA, UAS-IVS-myr-tdtomato / UAS-con-RNAi, UAS-Kek1, VT003280-LexA* | *R78H05-p65.AD, R22E04-GAL4.DBD, LexAop-IVS-jGCaMP7b / UAS-ten-a-RNAi, UAS-Fili, UAS-Ptp10D-RNAi* |
| **Figure 4** |  |  |  |  |
|  | Control male | *UAS-Dcr2 / y* | *VT003280-LexA, UAS-IVS-myr-tdtomato (BDSC 32222) / VT003280-LexA* | *R78H05-p65.AD, R22E04-GAL4.DBD, LexAop-IVS-jGCaMP7b (BDSC 80915) / +* |
|  | DA1-ORN rewired male | *UAS-Dcr2 / y* | *VT003280-LexA, UAS-IVS-myr-tdtomato / UAS-con-RNAi, UAS-Kek1, VT003280-LexA* | *R78H05-p65.AD, R22E04-GAL4.DBD, LexAop-IVS-jGCaMP7b / UAS-ten-a-RNAi, UAS-Fili, UAS-Ptp10D-RNAi* |
| **Figure 5** |  |  |  |  |
| b, c | VA1d-PN & wild type | *UAS-Dcr2, UAS-mCD8-GFP / w or y* | *R20D10-QF2<sup>10</sup>, QUAS-mtdTomato-HA / +* | *R78H05-p65.AD, R31F09-GAL4.DBD (BDSC 68759) / +* |
|  | VA1d-PN & CSP set 1 | *UAS-Dcr2, UAS-mCD8-GFP / w or y* | *R20D10-QF2, QUAS-mtdTomato-HA / UAS-con-RNAi, UAS-Kek1* | *R78H05-p65.AD, R31F09-GAL4.DBD / UAS-Ptp10D-RNAi, UAS-Sema-2b-RNAi (BDSC 28932)* |
|  | VA1d-PN & CSP set 2 | *UAS-Dcr2, UAS-mCD8-GFP / UAS-Kirre* | *R20D10-QF2, QUAS-mtdTomato-HA / UAS-Kek1* | *R78H05-p65.AD, R31F09-GAL4.DBD / UAS-Ptp10D-RNAi, UAS-Ten-m-RNAi (BDSC 29390), UAS-Klg<sup>17</sup>* |
|  | VA1d-PN & CSP set 3 | *UAS-Dcr2, UAS-mCD8-GFP / UAS-Ten-a (BDSC 41563)* | *R20D10-QF2, QUAS-mtdTomato-HA / +* | *R78H05-p65.AD, R31F09-GAL4.DBD / UAS-Sema-2b-RNAi* |
|  | VA1v-PN & wild type | *UAS-Dcr2 / w or y* | *VT003280-LexA, lexAop-rCD2::RFP-p10.UAS-mCD8::GFP-p10 / +* | *R78H05-p65.AD, R31F09-GAL4.DBD / +* |
|  | VA1v-PN & CSP set 1 | *UAS-Dcr2 / w or y* | *VT003280-LexA, lexAop-rCD2::RFP-p10.UAS-mCD8::GFP-p10 / UAS-con-RNAi, UAS-Kek1* | *R78H05-p65.AD, R31F09-GAL4.DBD / UAS-Ptp10D-RNAi, UAS-Sema-2b-RNAi* |
|  | VA1v-PN & CSP set 2 | *UAS-Dcr2 / UAS-Kirre* | *VT003280-LexA, lexAop-rCD2::RFP-p10.UAS-mCD8::GFP-p10 / UAS-Kek1* | *R78H05-p65.AD, R31F09-GAL4.DBD / UAS-Ptp10D-RNAi, UAS-Ten-m-RNAi, UAS-Klg* |
|  | VA1v-PN & CSP set 3 | *UAS-Dcr2 / UAS-Ten-a* | *VT003280-LexA, lexAop-rCD2::RFP-p10.UAS-mCD8::GFP-p10 / +* | *R78H05-p65.AD, R31F09-GAL4.DBD / UAS-Sema-2b-RNAi* |
|  | DC3-PN & wild type | *UAS-Dcr2, UAS-mCD8-GFP / w or y* | *R82E01-QF2<sup>10</sup>, QUAS-mtdTomato-HA / +* | *R78H05-p65.AD, R31F09-GAL4.DBD / +* |
|  | DC3-PN & CSP set 1 | *UAS-Dcr2, UAS-mCD8-GFP / w or y* | *R82E01-QF2, QUAS-mtdTomato-HA / UAS-con-RNAi, UAS-Kek1* | *R78H05-p65.AD, R31F09-GAL4.DBD / UAS-Ptp10D-RNAi, UAS-Sema-2b-RNAi* |
|  | DC3-PN & CSP set 2 | *UAS-Dcr2, UAS-mCD8-GFP / UAS-Kirre* | *R82E01-QF2, QUAS-mtdTomato-HA / UAS-Kek1* | *R78H05-p65.AD, R31F09-GAL4.DBD / UAS-Ptp10D-RNAi, UAS-Ten-m-RNAi, UAS-Klg* |
|  | DC3-PN & CSP set 3 | *UAS-Dcr2, UAS-mCD8-GFP / UAS-Ten-a* | *R82E01-QF2, QUAS-mtdTomato-HA / +* | *R78H05-p65.AD, R31F09-GAL4.DBD / UAS-Sema-2b-RNAi* |
|  | DL3-PN & wild type | *UAS-Dcr2 / w or y* | *R37H08-LexA (BDSC 52766), lexAop-rCD2::RFP-p10.UAS-mCD8::GFP-p10 / +* | *R78H05-p65.AD, R31F09-GAL4.DBD / +* |
|  | DL3-PN & CSP set 1 | *UAS-Dcr2 / w or y* | *R37H08-LexA, lexAop-rCD2::RFP-p10.UAS-mCD8::GFP-p10 / UAS-con-RNAi, UAS-Kek1* | *R78H05-p65.AD, R31F09-GAL4.DBD / UAS-Ptp10D-RNAi, UAS-Sema-2b-RNAi* |
|  | DL3-PN & CSP set 2 | *UAS-Dcr2 / UAS-Kirre* | *R37H08-LexA, lexAop-rCD2::RFP-p10.UAS-mCD8::GFP-p10 / UAS-Kek1* | *R78H05-p65.AD, R31F09-GAL4.DBD / UAS-Ptp10D-RNAi, UAS-Ten-m-RNAi, UAS-Klg* |
|  | DL3-PN & CSP set 3 | *UAS-Dcr2 / UAS-Ten-a* | *R37H08-LexA, lexAop-rCD2::RFP-p10.UAS-mCD8::GFP-p10 / +* | *R78H05-p65.AD, R31F09-GAL4.DBD / UAS-Sema-2b-RNAi* |
| d, e, j | controls | *UAS-Dcr2 / w* | *VT003280-LexA, UAS-IVS-myr-tdtomato / VT003280-LexA* | *R78H05-p65.AD, R31F09-GAL4.DBD, LexAop-IVS-jGCaMP7b / +* |
|  | VA1d-ORN to VA1v-PN rewired | *UAS-Dcr2 / w* | *VT003280-LexA, UAS-IVS-myr-tdtomato / VT003280-LexA, UAS-con-RNAi, UAS-Kek1* | *R78H05-p65.AD, R31F09-GAL4.DBD, LexAop-IVS-jGCaMP7b / UAS-Ptp10D-RNAi, UAS-Sema-2b-RNAi* |
| f, g, k | controls | *UAS-Dcr2, QUAS-GCaMP8m (this work) / +* | *R82E01-QF2, UAS-IVS-myr-tdtomato / R82E01-QF2* | *R78H05-p65.AD, R31F09-GAL4.DBD, QUAS-GCaMP8m (this work) / +* |
|  | VA1d-ORN to DC3-PN rewired | *UAS-Dcr2, QUAS-GCaMP8m / UAS-Kirre* | *R82E01-QF2, UAS-IVS-myr-tdtomato / UAS-Kek1, R82E01-QF2* | *R78H05-p65.AD, R31F09-GAL4.DBD, QUAS-GCaMP8m / UAS-Ptp10D-RNAi, UAS-Ten-m-RNAi, UAS-Klg* |
| h, i, l | controls | *UAS-Dcr2 / w* | *R37H08-LexA, UAS-IVS-myr-tdtomato / R37H08-LexA* | *R78H05-p65.AD, R31F09-GAL4.DBD, LexAop-IVS-jGCaMP7b / +* |
|  | VA1d-ORN to DL3-PN rewired | *UAS-Dcr2 / UAS-Ten-a* | *R37H08-LexA, UAS-IVS-myr-tdtomato / R37H08-LexA* | *R78H05-p65.AD, R31F09-GAL4.DBD, LexAop-IVS-jGCaMP7b / UAS-Sema-2b-RNAi* |
| **Extended Data Figure. 1** |  |  |  |  |
|  | DA1-ORN | *UAS-mCD8-GFP / w or y* | + | *R78H05-p65.AD, R22E04-GAL4.DBD / +* |
| **Extended Data Figure. 2** |  |  |  |  |
| c |  | *UAS-Dcr2 / w or y* | *Mz19-GAL4 (BDSC 34497), QUAS_mCD8_GFP (BDSC 30002), Or47b_rCD2, Or88a_mtdTomato (BDSC 64070) / +* | + |
| d |  | *UAS-Dcr2 / w or y* | *Mz19-GAL4, QUAS_mCD8_GFP, Or47b_rCD2, Or88a_mtdTomato / UAS-con-RNAi* | + |
| e |  | *UAS-Dcr2, UAS-mCD8-GFP / w or y* | *R20D10-QF2, QUAS-mtdTomato-HA / +* | *R78H05-p65.AD, R31F09-GAL4.DBD / +* |
| f |  | *UAS-Dcr2, UAS-mCD8-GFP / w or y* | *R20D10-QF2, QUAS-mtdTomato-HA / +* | *R78H05-p65.AD, R31F09-GAL4.DBD / UAS-Klg* |
| **Extended Data Figure. 5** |  |  |  |  |
|  | No *ptp10D* RNAi with DA1-PN colabeling | *UAS-Dcr2, UAS-mCD8-GFP / w or y* | *R24A10-QF2, QUAS-mtdTomato-HA / UAS-con-RNAi, UAS-Kek1* | *R78H05-p65.AD, R22E04-GAL4.DBD / UAS-Ten-a-RNAi, UAS-Fili* |
|  | No *Ten-a* RNAi with DA1-PN colabeling | *UAS-Dcr2, UAS-mCD8-GFP / w or y* | *R24A10-QF2, QUAS-mtdTomato-HA / UAS-con-RNAi, UAS-Kek1* | *R78H05-p65.AD, R22E04-GAL4.DBD / UAS-Fili, UAS-Ptp10D-RNAi* |
|  | No *con* RNAi with DA1-PN colabeling | *UAS-Dcr2, UAS-mCD8-GFP / w or y* | *R24A10-QF2, QUAS-mtdTomato-HA / UAS-Kek1* | *R78H05-p65.AD, R22E04-GAL4.DBD / UAS-Ten-a-RNAi, UAS-Fili, UAS-Ptp10D-RNAi* |
|  | No *Kek1* OE with DA1-PN colabeling | *UAS-Dcr2, UAS-mCD8-GFP / w or y* | *R24A10-QF2, QUAS-mtdTomato-HA / UAS-con-RNAi* | *R78H05-p65.AD, R22E04-GAL4.DBD / UAS-Ten-a-RNAi, UAS-Fili, UAS-Ptp10D-RNAi* |
|  | No Fili OE with DA1-PN colabeling | *UAS-Dcr2, UAS-mCD8-GFP / w or y* | *R24A10-QF2, QUAS-mtdTomato-HA / UAS-con-RNAi, UAS-Kek1* | *R78H05-p65.AD, R22E04-GAL4.DBD / UAS-Ten-a-RNAi, UAS-Ptp10D-RNAi* |
|  | No *ptp10D* RNAi with VA1v-PN colabeling | *UAS-Dcr2, UAS-mCD8-GFP / w or y* | *VT003280-LexA, lexAop-rCD2::RFP-p10.UAS-mCD8::GFP-p10 / UAS-con-RNAi, UAS-Kek1* | *R78H05-p65.AD, R22E04-GAL4.DBD / UAS-Ten-a-RNAi, UAS-Fili* |
|  | No *Ten-a* RNAi with VA1v-PN colabeling | *UAS-Dcr2, UAS-mCD8-GFP / w or y* | *VT003280-LexA, lexAop-rCD2::RFP-p10.UAS-mCD8::GFP-p10 / UAS-con-RNAi, UAS-Kek1* | *R78H05-p65.AD, R22E04-GAL4.DBD / UAS-Fili, UAS-Ptp10D-RNAi* |
|  | No *con* RNAi with VA1v-PN colabeling | *UAS-Dcr2, UAS-mCD8-GFP / w or y* | *VT003280-LexA, lexAop-rCD2::RFP-p10.UAS-mCD8::GFP-p10 / UAS-Kek1* | *R78H05-p65.AD, R22E04-GAL4.DBD / UAS-Ten-a-RNAi, UAS-Fili, UAS-Ptp10D-RNAi* |
|  | No *Kek1* OE with VA1v-PN colabeling | *UAS-Dcr2, UAS-mCD8-GFP / w or y* | *VT003280-LexA, lexAop-rCD2::RFP-p10.UAS-mCD8::GFP-p10 / UAS-con-RNAi* | *R78H05-p65.AD, R22E04-GAL4.DBD / UAS-Ten-a-RNAi, UAS-Fili, UAS-Ptp10D-RNAi* |
|  | No Fili OE with VA1v-PN colabeling | *UAS-Dcr2, UAS-mCD8-GFP / w or y* | *VT003280-LexA, lexAop-rCD2::RFP-p10.UAS-mCD8::GFP-p10 / UAS-con-RNAi, UAS-Kek1* | *R78H05-p65.AD, R22E04-GAL4.DBD / UAS-Ten-a-RNAi, UAS-Ptp10D-RNAi* |
|  | 5-CSP manipulation | *Same as Fig. 2c* |  |  |
| **Extended Data Figure. 6** |  |  |  |  |
|  | rewired male with DA1-PN colabeling | *UAS-Dcr2, UAS-mCD8-GFP / y* | *R24A10-QF2, QUAS-mtdTomato-HA / UAS-con-RNAi, UAS-Kek1* | *R78H05-p65.AD, R22E04-GAL4.DBD / UAS-Ten-a-RNAi, UAS-Fili, UAS-Ptp10D-RNAi* |
|  | rewired male with VA1v-PN colabeling | *UAS-Dcr2, UAS-mCD8-GFP / y* | *VT003280-LexA, lexAop-rCD2::RFP-p10.UAS-mCD8::GFP-p10 / UAS-con-RNAi, UAS-Kek1* | *R78H05-p65.AD, R22E04-GAL4.DBD / UAS-Ten-a-RNAi, UAS-Fili, UAS-Ptp10D-RNAi* |
|  | rewired female with DA1-PN colabeling | *UAS-Dcr2, UAS-mCD8-GFP / w* | *R24A10-QF2, QUAS-mtdTomato-HA / UAS-con-RNAi, UAS-Kek1* | *R78H05-p65.AD, R22E04-GAL4.DBD / UAS-Ten-a-RNAi, UAS-Fili, UAS-Ptp10D-RNAi* |
|  | rewired female with VA1v-PN colabeling | *UAS-Dcr2, UAS-mCD8-GFP / w* | *VT003280-LexA, lexAop-rCD2::RFP-p10.UAS-mCD8::GFP-p10 / UAS-con-RNAi, UAS-Kek1* | *R78H05-p65.AD, R22E04-GAL4.DBD / UAS-Ten-a-RNAi, UAS-Fili, UAS-Ptp10D-RNAi* |
|  | DA1-ORN rewired with VA1d-ORN colabeling | *UAS-Dcr2, UAS-mCD8-GFP / w* | *Mz19-QF, QUAS-mCD8GFP, Or47b-rCD2, Or88a-mtdT<sup>41</sup> / UAS-con-RNAi, UAS-Kek1* | *R78H05-p65.AD, R22E04-GAL4.DBD / UAS-Ten-a-RNAi, UAS-Fili, UAS-Ptp10D-RNAi* |
|  | Control for VA1d-ORN colabeling | *UAS-Dcr2, UAS-mCD8-GFP / w* | *Mz19-QF, QUAS-mCD8GFP, Or47b-rCD2, Or88a-mtdT / +* | *R78H05-p65.AD, R22E04-GAL4.DBD / +* |
|  | DA1-ORN rewired with VA2-ORN colabeling | *UAS-Dcr2, UAS-mCD8-GFP / w* | *Or92a-rCD2, Or98a-mCD8-GFP<sup>9</sup> / UAS-con-RNAi, UAS-Kek1* | *R78H05-p65.AD, R22E04-GAL4.DBD / UAS-Ten-a-RNAi, UAS-Fili, UAS-Ptp10D-RNAi* |
|  | Control for VA2-ORN colabeling | *UAS-Dcr2, UAS-mCD8-GFP / w* | *Or92a-rCD2, Or98a-mCD8-GFP / +* | *R78H05-p65.AD, R22E04-GAL4.DBD / +* |
| **Extended Data Figure. 7** | Same as **Fig. 3** |  |  |  |
| **Extended Data Figure. 8** |  |  |  |  |
| b, c | Control male | *UAS-Dcr2 / y* | *VT003280-LexA, UAS-IVS-myr-tdtomato (BDSC 32222) / VT003280-LexA* | *R78H05-p65.AD, R22E04-GAL4.DBD, LexAop-IVS-jGCaMP7b (BDSC 80915) / +* |
|  | DA1-ORN rewired male | *UAS-Dcr2 / y* | *VT003280-LexA, UAS-IVS-myr-tdtomato / UAS-con-RNAi, UAS-Kek1, VT003280-LexA* | *R78H05-p65.AD, R22E04-GAL4.DBD, LexAop-IVS-jGCaMP7b / UAS-Ten-a-RNAi, UAS-Fili, UAS-Ptp10D-RNAi* |
|  | female | *Canton-S (BDSC 64349)* |  |  |
| e-j | rewired male with DA1-ORN expressing Kir2.1 | *UAS-Dcr2 / w* | *UAS-Kir2.1-T2A-tdtomato<sup>42</sup> / UAS-con-RNAi, UAS-Kek1, VT003280-LexA* | *R78H05-p65.AD, R22E04-GAL4.DBD, LexAop-IVS-jGCaMP7b / UAS-Ten-a-RNAi, UAS-Fili, UAS-Ptp10D-RNAi* |
|  | rewired male with DA1-ORN expressing csChrimson | *UAS-Dcr2 / w* | *UAS-csChrimson-mCherry<sup>43</sup> / UAS-con-RNAi, UAS-Kek1, VT003280-LexA* | *R78H05-p65.AD, R22E04-GAL4.DBD, LexAop-IVS-jGCaMP7b / UAS-Ten-a-RNAi, UAS-Fili, UAS-Ptp10D-RNAi* |
|  | Control male for the Kir2.1 group | *UAS-Dcr2 / w* | *UAS-Kir2.1-T2A-tdtomato<sup>42</sup> / VT003280-LexA* | *R78H05-p65.AD, R22E04-GAL4.DBD, LexAop-IVS-jGCaMP7b / +* |
|  | Control male for the csChrimson group | *UAS-Dcr2 / w* | *UAS-csChrimson-mCherry<sup>43</sup> / VT003280-LexA* | *R78H05-p65.AD, R22E04-GAL4.DBD, LexAop-IVS-jGCaMP7b / +* |
| k-l | male with DA1-ORN expressing Kir2.1 | *UAS-Dcr2 / w* | *UAS-Kir2.1-T2A-tdtomato<sup>42</sup> / +* | *R78H05-p65.AD, R22E04-GAL4.DBD / +* |
|  | Control male | *UAS-Dcr2 / w* | *+ / +* | *R78H05-p65.AD, R22E04-GAL4.DBD / +* |
| **Extended Data Figure. 10** |  |  |  |  |
| a | VA1v-PN & CSP set 1 without trajectory manipulation | *UAS-Dcr2 / w or y* | *VT003280-LexA, lexAop-rCD2::RFP-p10.UAS-mCD8::GFP-p10 / UAS-Con-RNAi, UAS-Kek1* | *R78H05-p65.AD, R31F09-GAL4.DBD / UAS-Ptp10D-RNAi* |
| b | DL3-PN & CSP set 3 without trajectory manipulation | *UAS-Dcr2 / UAS-Ten-a* | *R37H08-LexA, lexAop-rCD2::RFP-p10.UAS-mCD8::GFP-p10 / +* | *R78H05-p65.AD, R31F09-GAL4.DBD / +* |
| c, d | Same as Fig. 5b, but without *Sema-2b* RNAi in groups that do not have trajectory manipulation |  |  |  |

## Supplementary Materials

**Video 1** Courtship activity of two virgin males: a control male and a DA1-ORNs→VA1v-PNs rewired male with a white dot on the thorax. The behavioral chamber has a diameter of 2 cm.

**Video 2** Courtship activity of two virgin males: a control male and a DA1-ORNs→VA1v-PNs rewired male with a white dot on the thorax. The behavioral chamber has a diameter of 5 cm. Sped up to x5 at approximately 18 second. The lighting is stronger than in Video 1 and the courtship behavior is in general more continuous and vigorous than when the lighting is dimmer as shown in Video 1.

**Video 3** Courtship activity of five virgin DA1-ORNs→VA1v-PNs males. The behavioral chamber has a diameter of 5 cm.

